# An associative transcriptomics study on rice bean (*Vigna umbellata*) provides new insights into genetic basis and candidate genes governing flowering, maturity and seed weight

**DOI:** 10.1101/2023.02.25.530014

**Authors:** Tanmaya Kumar Sahu, Sachin Kumar Verma, Gayacharan, Nagendra Pratap Singh, Dinesh Chandra Joshi, D. P. Wankhede, Mohar Singh, Rakesh Bhardwaj, Swarup Kumar Parida, Debasis Chattopadhyay, Gyanendra Pratap Singh, Amit Kumar Singh

## Abstract

Rice bean is an underrated legume with significant potential to support food and nutritional security worldwide, being a rich source of proteins, minerals, and essential fatty acids. Therefore, we considered three pivotal production traits of rice bean; flowering, maturity and seed weight, to identify associated candidate genes. One-hundred diverse genotypes out of 1800 evaluated rice bean accessions from the Indian National Genebank were considered for phenotypic data collection and genotyping by transcriptome sequencing approach. Association analysis involving various GWAS models was conducted to identify significant marker-trait associations. The results revealed association of 82 markers on 48 transcripts for flowering, 26 markers on 22 transcripts for maturity and 22 markers on 21 transcripts for seed weight. The annotation of associated transcripts unraveled the functional genes related to the considered traits. Among the significant candidate genes identified, HSC80, P-II PsbX, phospholipid-transporting-ATPase-9, pectin-acetylesterase-8 and E3-ubiquitin-protein-ligase-RHG1A were found associated with flowering. Further, associations of WRKY1 and DEAD-box-RH27 with seed weight, PIF3 and pentatricopeptide-repeat-containing-gene with maturity & seed weight and aldo-keto-reductase with flowering & maturity have been revealed. The present investigation provides insights into the genetic mechanisms governing economically-essential traits like flowering, maturity and seed weight that can be potentially utilized for rice bean improvement.

**Highlights:** The present investigation is the first associative transcriptomics approach implemented on ***Vigna umbellata***. Based on the marker-trait association analysis, candidate genes for flowering, maturity and seed weight traits are reported.

## Introduction

The rice bean [*Vigna umbellata* (Thunb.) Ohwi & Ohashi] is a comparatively short-duration, warm-season, annual, leguminous pulse crop. It is adaptive to diverse range of climatic conditions and can be grown in a wide range of soil types even in the poor quality soil. It is used as a vegetable, a minor food crop, a fodder crop and a green manure as well. Being a leguminous crop, rice bean has an advantageous role in mixed cropping and preventing soil erosion due to its ability to fix nitrogen in nutrient depleted soils. Rice bean is reportedly resistant to a wide range of biotic and abiotic stresses. Especially, it is a source of resistance against biotic stresses such as bruchids, yellow mosaic disease and bacterial leaf spots (Isemura *et al*., 2010). Among abiotic stresses, it is tolerant to some degree of water logging (de Carvalho and Veira, 1996), acid soils (Dwivedi, 1996), drought (NAS, 1979) and high temperatures.

Rice bean is important for the health of the dependent living beings as well as the soil. It has a great potential to overcome the food and nutritional deficiency across the globe (Pattanayak *et al*., 2019). In favorable climatic conditions, it produces large amounts of healthy animal fodder and superior quality of grains. It is a rich source of proteins, minerals, essential fatty acids and amino acids (Mohan and Janardhanan, 1994). Its amino acid composition is reported to be well balanced for human consumption (Chandel *et al*., 1978; Mohan and Janardhan, 1994; de Carvalho and Vieira, 1996). Though it is a beneficial, nutritious and low maintenance crop, a little research has been carried out on this crop till date. It has also remained as a neglected crop being cultivated on small areas in India, Nepal and parts of Southeast Asia (Dahiphale *et al*., 2017; Dhillon and Tanwar, 2018; Tian *et al*., 2013). Due to this negligence, it suffers with wild and disadvantageous traits such as indeterminate growth habit, asynchronous and late maturity, pod shattering and antinutritional compounds (Pattanayak *et al*., 2019). However, the level of enzyme inhibitors, anti-nutritive or toxic factors are low in rice bean as compared to other legumes (Smil, 1997).

Rice bean is a diploid crop with 11 pairs of chromosomes (2n=22) and has an estimated genome size of ~525.60 Mb (Guan *et al*., 2022). Availability of inadequate genomic information on rice bean has made the investigations on this crop challenging. A few genetic maps of rice bean have been constructed and utilized to localize genes for quantitative traits seed weight and several domestication related traits (Somta *et al*., 2006; Isemura *et al*., 2010). However, these maps do not provide high resolution mapping of targeted traits because they were based on small number of SSR markers derived from the related legume species (Isemura *et al*., 2010). Very recently in September 2022, one genome assembly of rice bean at chromosomal level has been released by Chinese Academy of Agricultural Sciences, Beijing, China (Guan *et al*., 2022) with the genome size of 475.64LMb that is 90.49% of the estimated genome size. However, another assembly by International Centre for Genetic Engineering and Biotechnology, New Delhi, India is available at scaffold level (Kaul *et al*., 2022) having the assembled genome size of 414 Mb. They estimated a total of 31276 highly confidential genes from 15521 scaffolds. Availability of these two assemblies opened an avenue for annotating markers and the corresponding candidate genes for important traits revealed through genome or transcriptome wide association analysis.

The cultivars of rice bean are highly photoperiod sensitive, therefore when the crop is grown in subtropical areas, vegetative growth continues for longer duration and crop flowers very late. Further, the rice bean crop improvement and its cultivation faces a challenge from other *Vigna* group of crops of similar nature, but with superior agronomic traits. Yield potential of rice bean crop in its present form is estimated to be 1200 kg/ha (Gautam *et al*., 2007), which is comparatively lesser than green gram, blackgram and cowpea. Similarly, indeterminate growth habit and pod shattering in rice bean makes the crop unsuitable for large scale or mechanized farming, and primarily grown as intercrop (Gautam *et al*., 2007). Owing to such constraints and the competition from other crops, rice bean cultivated area has gradually decreased, and even is being discontinued from traditional cultivation areas. However, recent trend towards food crop diversification amid climate change and health awareness of the society, rice bean crop is recognized as one of the most potential legume crops to meet out the current and future needs. Therefore, in this study we have selected three most important productivity related traits i.e., days to flowering, days to maturity after crop sowing and seed size to understand their genetics. Understanding the underlying genetic mechanism governing these traits will help in developing rice bean genotypes with early flowering and early maturity with bolder seeds, which will enhance the large scale area expansion under rice bean crop.

Genome wide association analysis involving genotypic and phenotypic data has empowered efficient detection of significant markers associated with the trait of interest (Zhao *et al*., 2011, Xiao *et al*., 2017). If interest lies in the expressed part of the genome, variant calling from transcriptome sequencing can be adopted, which is comparatively easier, cost effective and time efficient than whole genome sequencing (Hong *et al*., 2022, Jehl *et al*., 2021). Further, transcriptome-based variant calling is expected to capture the variants in the parts of the transcripts modified during to post-transcriptional modifications and RNA editing. Therefore, we attempted here transcriptome-based variant calling along with association analysis to unravel the putative candidate genes for important productivity related traits like, flowering, maturity and seed weight of Rice bean. The comprehensive associative transcriptome analysis on rice bean explored here is expected to enlighten the stakeholders on the implementation of transcriptome-based variant calling to identify putative candidate genes for economically important traits in other crops too.

## Materials and methods

### Plant material

A diverse set of 100 accessions of rice bean was selected based on the phenotypic characteristics of 1800 accessions which were acquired from Indian National Genebank (http://pgrportal.nbpgr.ernet.in/) and characterized for important agro-morphological traits during 2018 and 2019. We also tried to make selection of these accessions in such a way that each rice bean growing geographical region of India is represented (Fig. 1). To avoid any admixture single plant selection representative to the accession population was made. Based on the phenotypic characteristic information generated from entire set of rice bean accessions, 100 accessions were selected for this study having good range of variation for flowering and maturity period, and seed size. The selected 100 accessions were further grown during rainy season of years 2020 and 2021 at the Experimental Farm of ICAR-National Bureau of Plant Genetic Resources, Issapur (28° 34’25”N, 76°50’41”E, 215 m msl) and Experimental farm Hawalbagh, ICAR-Vivekananda Institute of Hill Agriculture, Almora (79.39° E longitude and 25.35° N latitude, mean rainfall-1000 mm and 1250 m above msl) following augmented block design. Each accession was showed in paired rows of 2 m length and row-to-row distance 60 cm and plant to plant distance 30 cm. The standard rice bean growing practices were followed to obtain ideal phenotypic expression of genotypes. The detailed passport information about these accessions such as date of collection, site of origin and cultivar name are given in the Table S1. The phenotypic data for 100 accessions including “days to 50% flowering”, “days to 80% maturity” and “100 seed weight” were recorded for all the accessions.

**Fig. 1.**
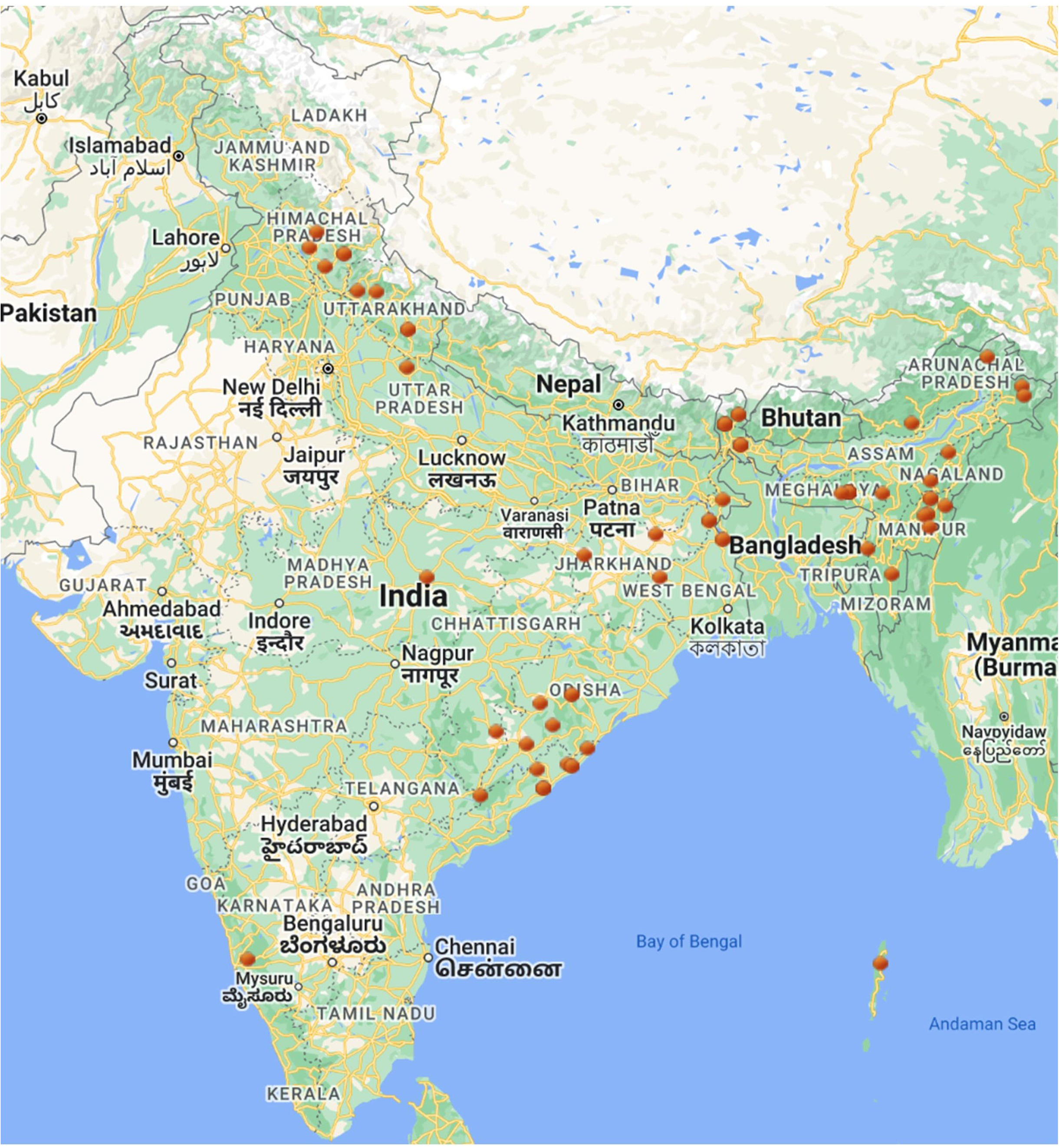
Geographical distribution of 100 rice bean accessions considered under this study

### Data acquisition and processing

#### Phenotypic data analysis

The phenotypic data collected from two locations in two consecutive years was analyzed and descriptive statistics were obtained. To detect the significant variations in the datasets analysis of variance (ANOVA) for each trait was carried out in the interface of SPSS version 15. The frequency distributions for each trait for all datasets were analyzed. Further, trait-wise comparative density plots were generated using *sm* library (Bowman and Azzalini, 2021) of R.

#### Genotypic data generation

These accessions were further genotyped by transcriptome sequencing followed by variant calling. The transcriptome sequencing was carried out in Illumina Hiseq6000 platform. Cleaned Illumina short reads were obtained by fastp v0.12 from raw sequencing data. Single-nucleotide polymorphisms (SNPs) were identified using GATK version v4.1 by mapping cleaned short reads against the Rice bean transcriptome reference VRB3 using BWA-mem v0.7. Duplicate reads were marked by Picard tools. Genomic variants were identified using GATK HaplotypeCaller, and a joint variant call set was generated using GATK Genotype GVCFs. Subsequently, SNP variants were selected and filtered to retain high-quality SNPs.

#### Pre-processing of genotypic data

The genotypic data was pre-processed in the interface of TASSEL software (Trait Analysis by Association, Evolution and Linkage version 5. 2. 85; Bradbury *et al*., 2007). Initially, the markers were filtered out based on MAF (>5%), missing data (<20%) and heterozygosity (<50%). As rice bean is a cross pollinated crop the heterozygosity filter of 50% was implemented. Further, indels, triallelic markers and markers with minor state were removed. The genotypes were also examined for >30% missing data and > 50% heterozygosity.

### Population structure and diversity

To estimate the number of populations, the genotypic data was subjected to principal component analysis (PCA) and population structure analysis. The PCA was carried out in the graphical user interface of TASSEL on the filtered set of markers and the first three principal components were plotted to analyze the population structure. To execute the admixture model of STRUCTURE v2.3.4 (Pritchard *et al*., 2000) with Bayesian Markov Chain Monte Carlo model (MCMC) simulation, the markers having high polymorphic information content (PIC) and genetic diversity (GD) were considered. The PIC and GD for each marker were computed based on the following formulae.

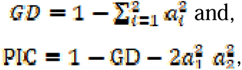

where, *a_i_* (i = 1, 2) is the frequency of i^th^ allele in the population. Here, the value of *i* varies from 1 to 2, because, we have considered only the bi-allelic markers. The markers with > 0.35 PIC were selected for STRUCTURE analysis. The structure software was executed with putative population (K=7), 25000 burning cycles, 50000 iterations of MCMC and 10 independent runs were executed for each K ranging from 2 to 7. The result of STRUCTURE was analyzed online with Structure Harvestor (Earl and vonHoldt, 2012) and the best value of K was selected based on highest delta K (ΔK). With the best assumed number of sub-populations, the structure simulation was rerun and the population membership file was used as the Q-matrix for the association analysis.

### Linkage Disequilibrium analysis

The linkage disequilibrium (LD) decay analysis was carried out to examine the genomic distance at which the LD starts decaying. Within this distance, the makers and genes are believed to have strong LD. Thus, the analysis of LD decay was carried out in the interface of TASSEL. As we have used transcriptome data, short distance LD has been examined by plotting the frequency of the squared allele of LD (r^2^) (Y-axis) against distance in base pair (X-axis) in the R command line for all pair-wise comparisons.

### Association analysis

The association analysis of markers with the considered traits was carried out using two single locus models (with two variations each) and six multi locus models. These models were implemented using TASSEL, GAPIT (Lipka *et al*., 2012) and mrMLM.GUI package (Wen *et al*., 2018).

#### Single locus models

Among the single locus models, generalized linear models (GLM; Price *et al*., 2006) and mixed linear model (MLM; Yu *et al*., 2006) have been used in this study. Here two variants of GLM have been used. In one variant, GLM (Q), the population membership information (Q matrix) was used as covariates whereas, in the other one, GLM (PCA), first three principal components (PC) were used as covariates to minimize the number of false positives. The MLM models are able to efficiently manage the population structure as well as relatedness within GWAS. Further, the incorporation of covariates in MLM controls the number of false positives (Gupta *et al*., 2019). Therefore, here we have also used two variants of MLM *i.e.,* MLM (Q+K) and MLM (PCA+K) using both Q-matrix and first three PCs respectively. However, in both the variants the kinship matrix (K) used, was generated based on identity by state analysis. The Q-matrix was generated by using STRUCTURE software, whereas the kinship matrix and principal components were generated in the TASSEL interface. All the four methods were implemented on the processed genotypic and phenotypic data of rice bean in the interface of TASSEL.

#### Multi locus models

Six multi-locus models viz., multi-locus random-SNP-effect MLM (mrMLM), fast multi-locus random-SNP-effect MLM (FASTmrMLM), fast multi-locus random-SNP-effect EMMA(FASTmrEMMA), Iterative Sure Independence Screening EM-Bayesian LASSO (ISIS EMBLASSO), fixed and random model circulating probability unification (FarmCPU) and Bayesian-information and linkage-disequilibrium iteratively nested keyway (BLINK) have been used in the study to analyze the marker-trait associations. The mrMLM being a multi-locus model avoids Bonferroni correction for multiple tests (Wang *et al*., 2016). FASTmrMLM is computationally faster and has considerable reliability as well as accuracy (Tamba and Zhang, 2018). FASTmrEMMA has also a low execution time and it estimates the QTN effect with low bias (Wen *et al*., 2018). ISIS EMBLASSO uses Expectation-Maximization (EM)-Bayesian least absolute shrinkage and selection operator (BLASSO) to estimate all the selected SNP effects for true quantitative trait nucleotide (QTN) detection (Tamba *et al*., 2017). In FarmCPU, the fixed effect model is used along with a random effect model iteratively to completely abolish the confounding effect (Liu *et al*., 2016). The random effect model in FarmCPU avoids over fitting and estimates the associated markers by employing them in defining the kinship. BLINK improves statistical power compared to FarmCPU, in addition to remarkably reducing computing time (Huang *et al*., 2019). Therefore, we have used these multi-locus models in our study. The FarmCPU and BLINK were implemented through GAPIT package where as all other multi-locus models were executed through mrMLM.GUI package.

### Marker selection

Markers have been selected based on different parameters for thresholds in different software packages. For the single locus models implemented through TASSEL, the markers were selected based on “–log10(p)” value > 5.99 after Bonferoni correction(Bland and Altman, 1995) i.e., 0.05/total number of markers. In the case of multi-locus models implemented through GAPIT, the markers were selected also based on “–log10(p)” value > 3.69(*i.e.,* 2 * 10^−4^; Zhang *et al*., 2019). However, in mrMLM a separate marker screening parameter *viz,* LOD score was used. Here markers were considered to be associated with a trait of interest if they have the LOD score >3 (Zhang *et al*., 2019). Though initial screening of the markers was based on the above defined parameters, the final screening was on the basis of their association predicted by at least two GWAS models considered under this study.

### Candidate gene identification

The transcripts on which the significant markers are located were subjected to a BLAST (Basic Local Alignment Search Tool) search for identifying the corresponding genes. To perform the BLAST search the transcript sequences were aligned to a local protein sequence database using blastx program of the offline BLAST. For creating a local database, the protein sequences of *Vigna* genus, G*lycine max* and *Arabidopsis* genus was collected from NCBI. As the later two plant species are well annotated and *Glycine max* is the close relative of rice bean apart from other *Vigna* members, the proteins sequences of these two plants were included to develop the local BLAST database. Further, all the transcripts were subjected to BLAST2GO of OmicsBox tool (https://www.biobam.com/omicsbox/) for annotation with gene ontology (GO) terms.

### Chromosomal localization of associated markers

To unravel the putative chromosomal location of markers as well as candidate genes, 50 bp left and 50 bp right flanking to the markers were extracted using a developed R-script and the resulting 101 bp fragments were subjected to the offline blastn program against the recently released fully sequenced genome of *V. umbellata* cultivar FF25(Guan *et al*., 2022). The exact marker locations were identified from the ungapped alignment of SNP sequences and chromosomal sequences obtained through blastn program. A chromosomal map of associated markers was created based on the newly released fully sequenced genome of *V. umbellata* in the interface of MapChart (Voorrips, 2002).

### Chromosomal localization of associated transcripts and synteny

The full length transcripts were subjected to blastn locally, against the genomes of *Vigna radiata, Vigna angularis, Vigna mungo, Vigna unguiculata* and the newly released genome of *V. umbellata* to identify their corresponding chromosomal locations. The chromosomal coordinates of the transcripts on *Vigna* genomes were used to carry out a synteny analysis using ShinyCircos software (Yu *et al*., 2017) to discover the synteny between these *Vigna* genomes with respect to the associated transcripts of *V. umbellata*.

### Expression analysis of associated transcripts

The expression of genes related to flowering, maturity and seed weight represented by the associated transcripts was checked in transcriptome data of inflorescence and developing seed tissues. The RNA sequencing reads of rice bean sample at young inflorescence stage (SRR5764826), 10 days post-anthesis stage (SRR16122602) and 5 days post-anthesis stage (SRR16122607) were downloaded. These reads were processed using FastQC(version 0.11.9) (Andrews, 2010), Trimomatic(version 0.40) (Bolger *et al*., 2014), bwa(Li and Durbin, 2009) and samtools (Li *et al*., 2009) for quality check, trimming, mapping and obtaining FPKM (fragments per kilobase of exon per million mapped fragments) values respectively. A heatmap of expression pattern was generated using an R script with heatmap() function.

## Results

### Analyzed phenotypic data

The phenotypic data for two years (2020 and 2021) from two locations (Delhi and Almora) for 100 selected accessions were analyzed and the data for some traits were found to be significantly different from each other either location-wise or year-wise (Fig. 2). The distribution of trait data for all the datasets is shown in the Fig. 3. The minimum, maximum and mean values of Almora datasets were found within a short range whereas in case of Delhi datasets these values are observed within a long range probably due to unexpectedly changing climatic conditions at Delhi. Further, the density plot for “days to 50% flowering” reveals similar distribution of Delhi datasets whereas the density plot for “100 seed weight” shows similar distribution of Almora datasets. However, for maturity data points of all the datasets were found to be equivalently distributed.

**Fig. 2.**
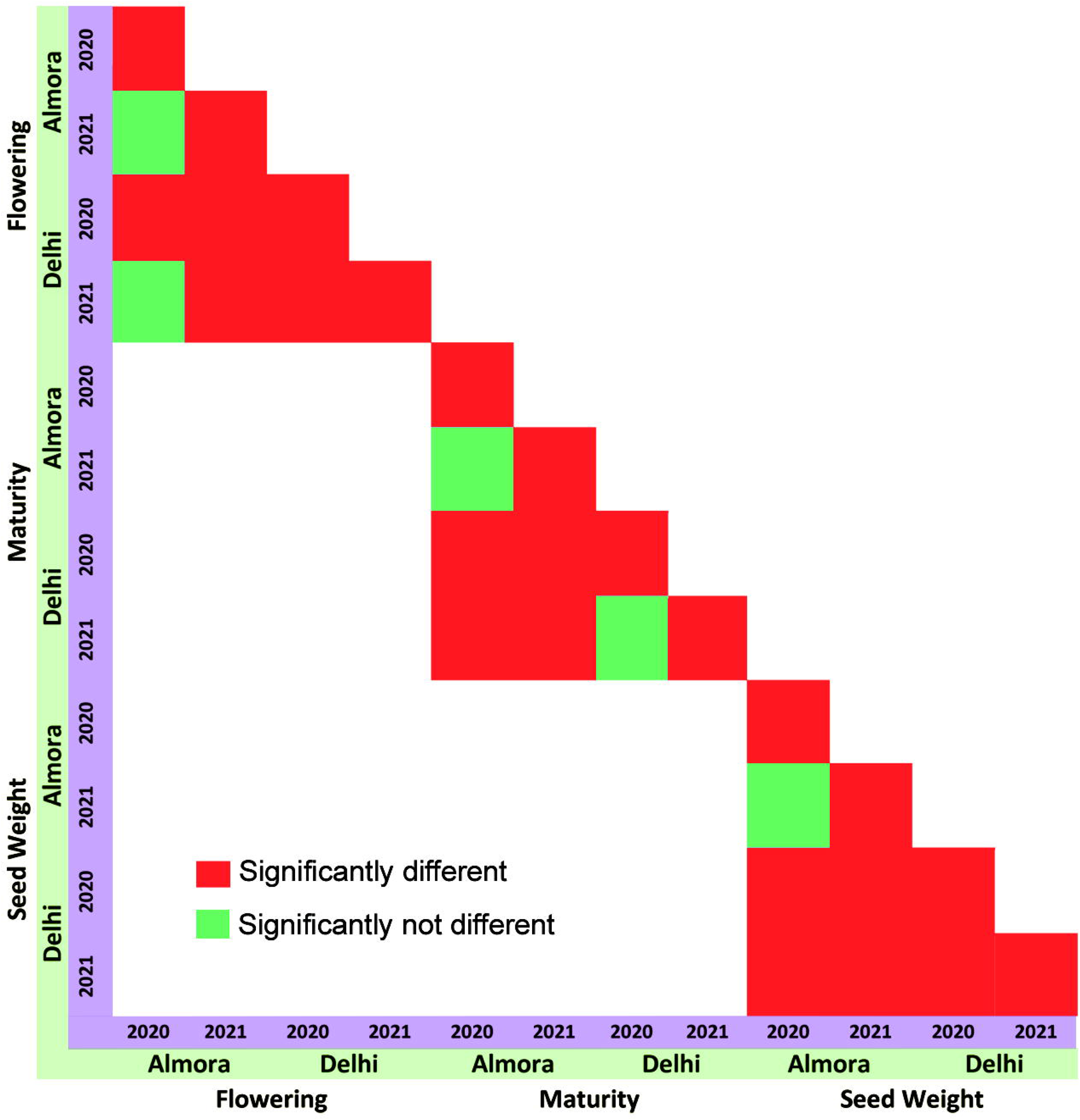
Analysis of significant difference in the phenotypic data of two years (2020-2021) from two locations (Delhi and Almora) based on ANOVA.

**Fig. 3.**
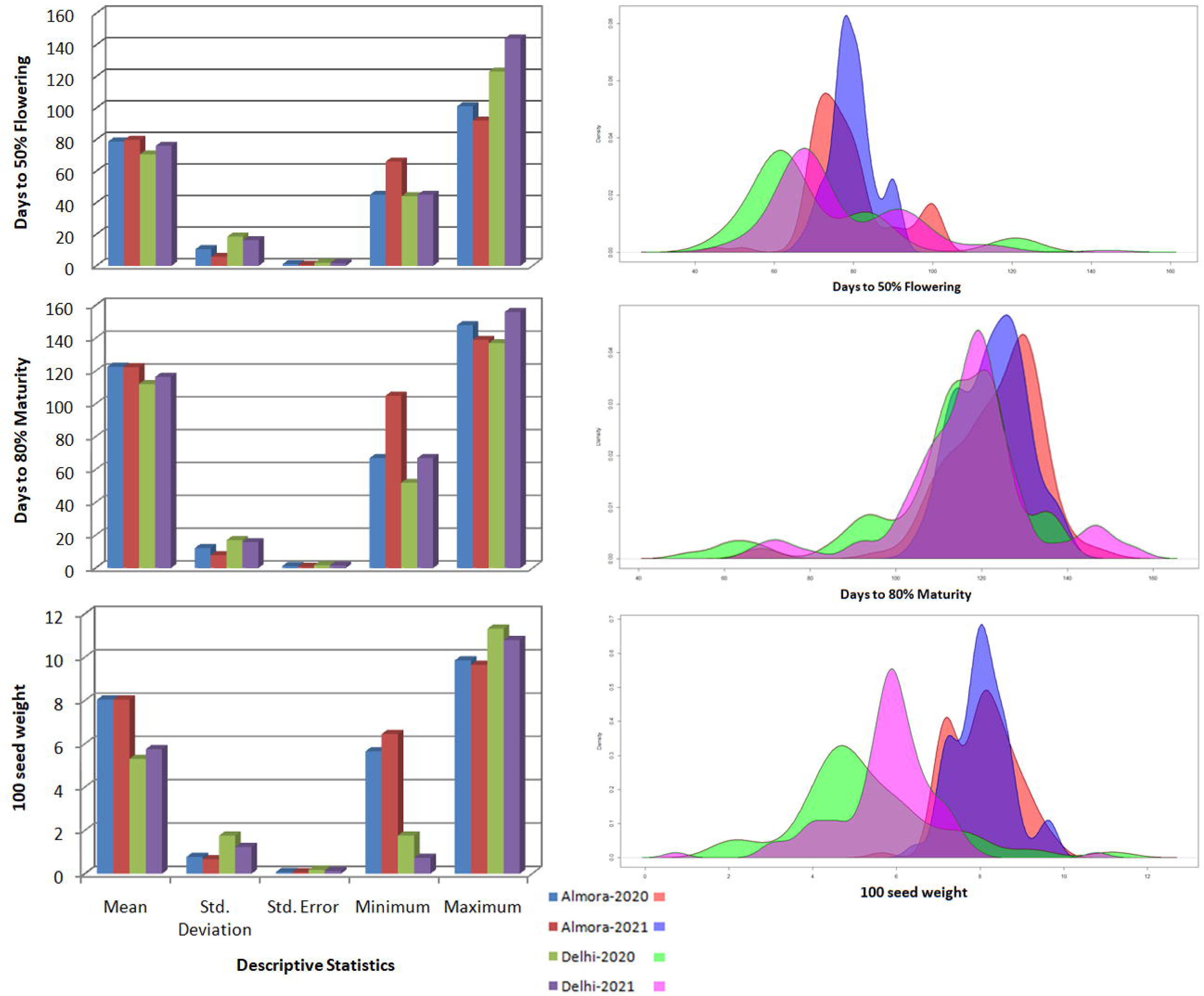
The distribution of phenotypic data under each trait for all the datasets showing the descriptive statistics and density plots.

### Processed genotypic data

The initial filtration of sequencing reads before variant calling generated cleaned short reads. The number of bases and reads before and after filtration is given in Table S2. The pre-processing of genotypic data with marker filtration parameters like indels, non bi-allelic markers, minor SNP states, minor allele frequency (<5%), missing genotype (>20%), heterozygosity(<50%) resulted 49271 markers. Further, the implementation of genotype filtration parameters like missing genotype (>30%) and heterozygosity (>50%) retained all the genotypes as all have missing data < 30% and heterozygosity < 50%. The final processed genotypic data contained 49271 markers for 100 genotypes for which the phenotypic data contained observations for 50% flowering, 80% maturity and 100 seed weight.

### Population structure

As simulation through STRUCTURE software takes high computational time with high number of markers, the structure simulation was executed with 5416 markers having PIC > 0.35. The calculated GD values varied from 0.0582 to 0.5 with an average of 0.269 whereas the PIC values varied from 0.0565 to 0.375 with an average of 0.2253 (Fig. 4). The markers having the PIC above 0.35 were found to have the GD above 0.45. Therefore, the STRUCTRE software was executed with the markers having high PIC and GD that is expected to correctly determine the number of populations. The result of STRUCTURE-MCMC simulation with K=1-7, when analyzed with Structure Harvestor online, revealed K=3 as the best value of K with highest ΔK (Fig. 5). Thus, the population structure estimated through admixture model of STRUCTRE-MCMC simulations revealed three putative populations (Fig. 6A). Further, three putative populations have also been revealed through the genotypic cluster (Fig. 6B) built in the Tassel interface by Neighbour Joining method and by plotting the first three principal components (Fig. 6C). Though the genotypic clustering reveals 3 distinct populations split from the root node, STRUCTRE showed a few admixed individuals in three populations. Large number of pure individuals was observed in the largest cluster (Population-1), followed by a smaller cluster (Population-2) than Population-1 and the smallest cluster (Population-3). Although Population-2 is larger than Population-3, the number of pure individuals in these populations are nearly equal. These three putative populations when matched with the passport information of the cultivars (Table S1), Population-1 was found to contain mostly the individuals of eastern and north-eastern regions of India and Population-2 was observed to have mostly the north Indian cultivars. The Population-3 was noticed to contain mixed individuals. However, few exotic cultivars were also noticed to fall within the Populations-1 & 2. The distribution of Indian cultivars is given in the Fig. 1.

**Fig. 4.**
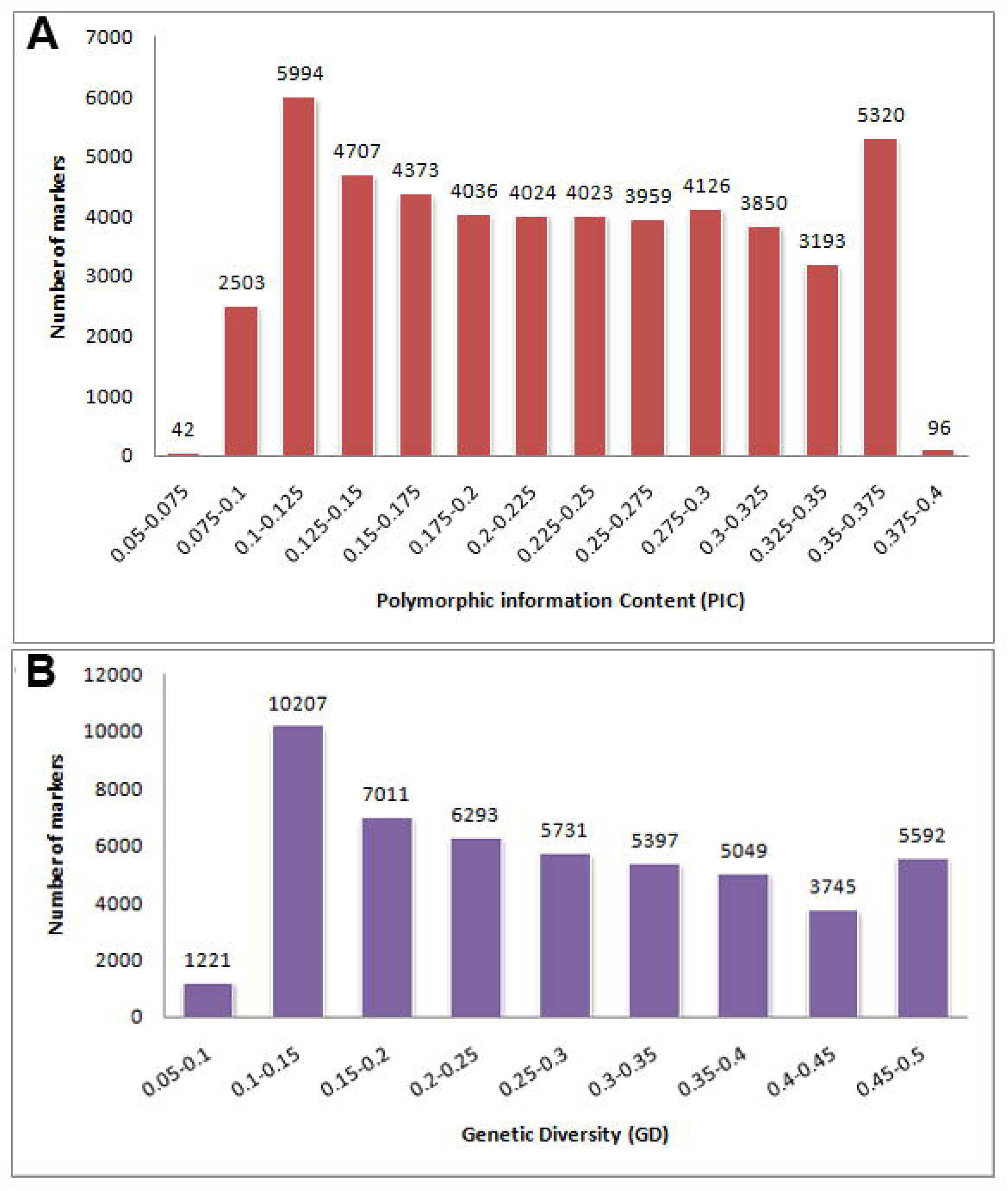
The distribution of genetic diversity (GD) and polymorphic information content (PIC) values calculated from the genotypic data.

**Fig. 5.**
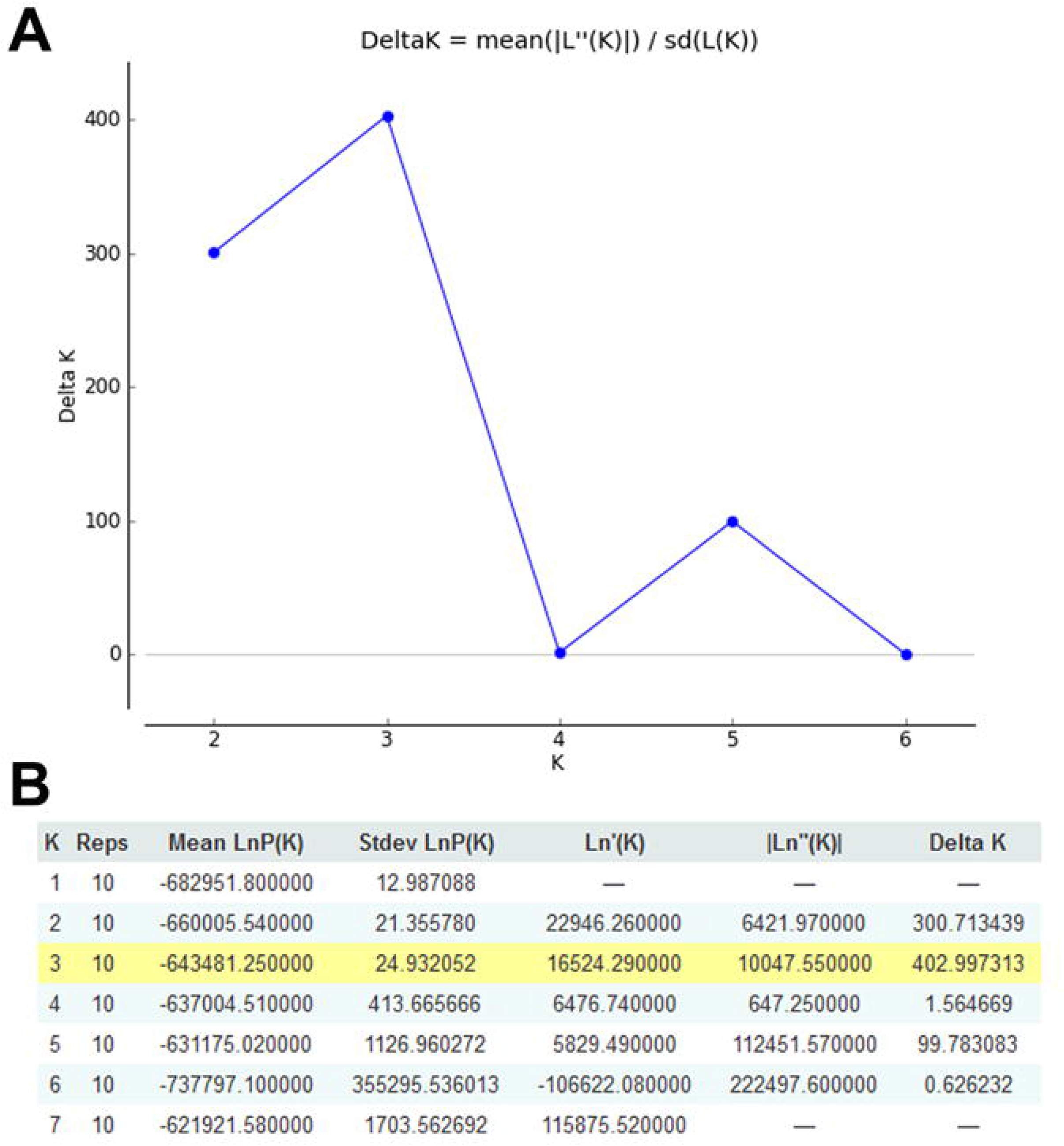
The graph showing the highest value of ΔK at K= 3(*i.e.,* three putative population). (A) Plot showing K in X-axis and ΔK in Y-axis. (B) Values of Mean LnP(K), Standard deviation LnP(K), Ln’(K), |Ln’’(K)| and ΔK for all values of K from 1 to 7.

**Fig. 6.**
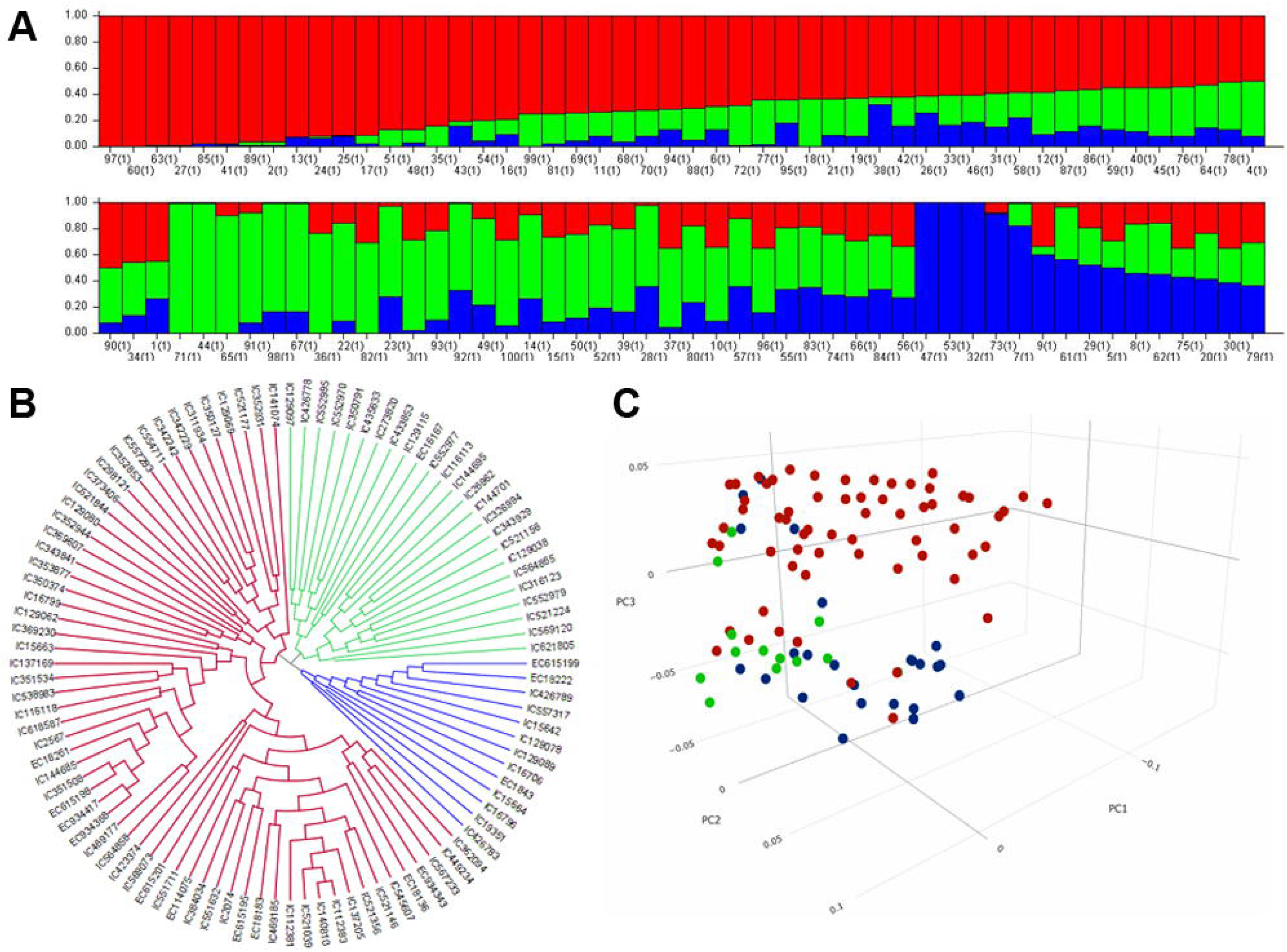
Three putative populations revealed from (A) the STRUCTRE-MCMC simulations, (B) the clustering based on Neighbour Joining method and (C) the plot of first three principal components.

### LD Decay

The decay in LD plotted by estimating the LD between all possible pair-wise marker-pairs presented in Fig. 7. The LD was observed to be decayed at a distance of 1.5Kb at the r^2^ cutoff of 2%. Further, the r^2^ was noticed to be on an average of 5% within a distance of 0.5Kb; however, it decayed rapidly from 4% to 2% within a distance of 0.5Kb to 2 Kb. After that the trend line seems to be in equilibrium up to a distance of 6Kb.

**Fig. 7.**
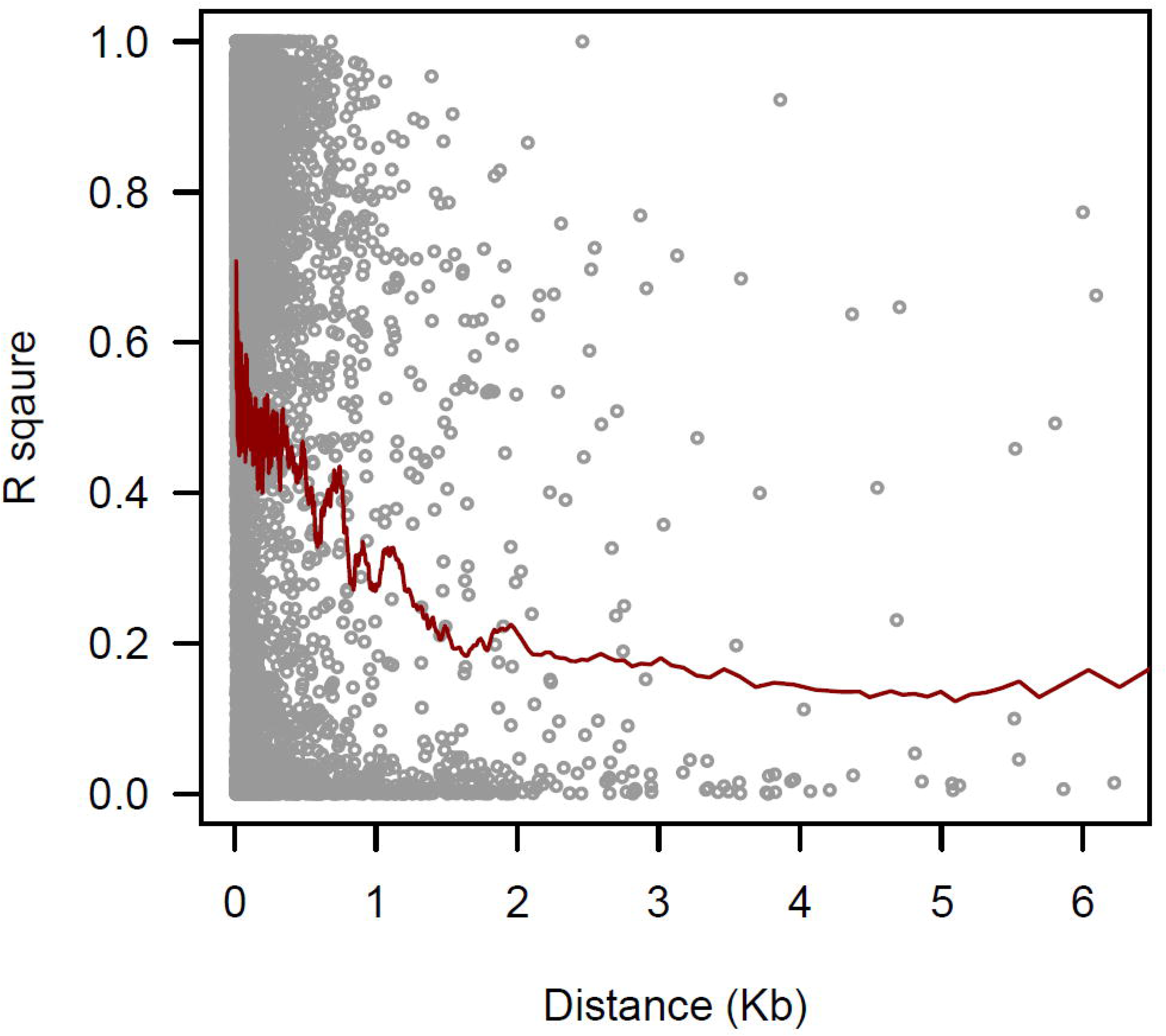
The decay in LD plotted by estimating the LD between all possible pair-wise markers. The LD decays at a distance of 1.5Kb at the r^2^ cutoff of 2%.

### Marker-trait association

#### Marker-trait association with Almora datasets

For Almora 2020, a good number of markers have been detected by both single and multi-locus models for all the considered traits (Table 1, Fig. 8). The markers considered here are predicted by at least two methods. With this dataset, 3 markers for flowering, 24 markers for maturity and 7 markers for seed weight have been predicted. However, 33 markers were identified to be associated with all the traits where one marker (SVUTC25856_1400) was found common for both maturity and seed weight traits. The marker, SVUTC06910_1648, predicted for maturity was predicted by 6 out of 10 GWAS models. Further, five markers at different positions (177, 1616, 1648, 1702, 1746) were found significant on one transcript VUTC06910 which are predicted to be associated with maturity trait. The remaining markers in this dataset were predicted on different transcripts.

**Fig. 8.**
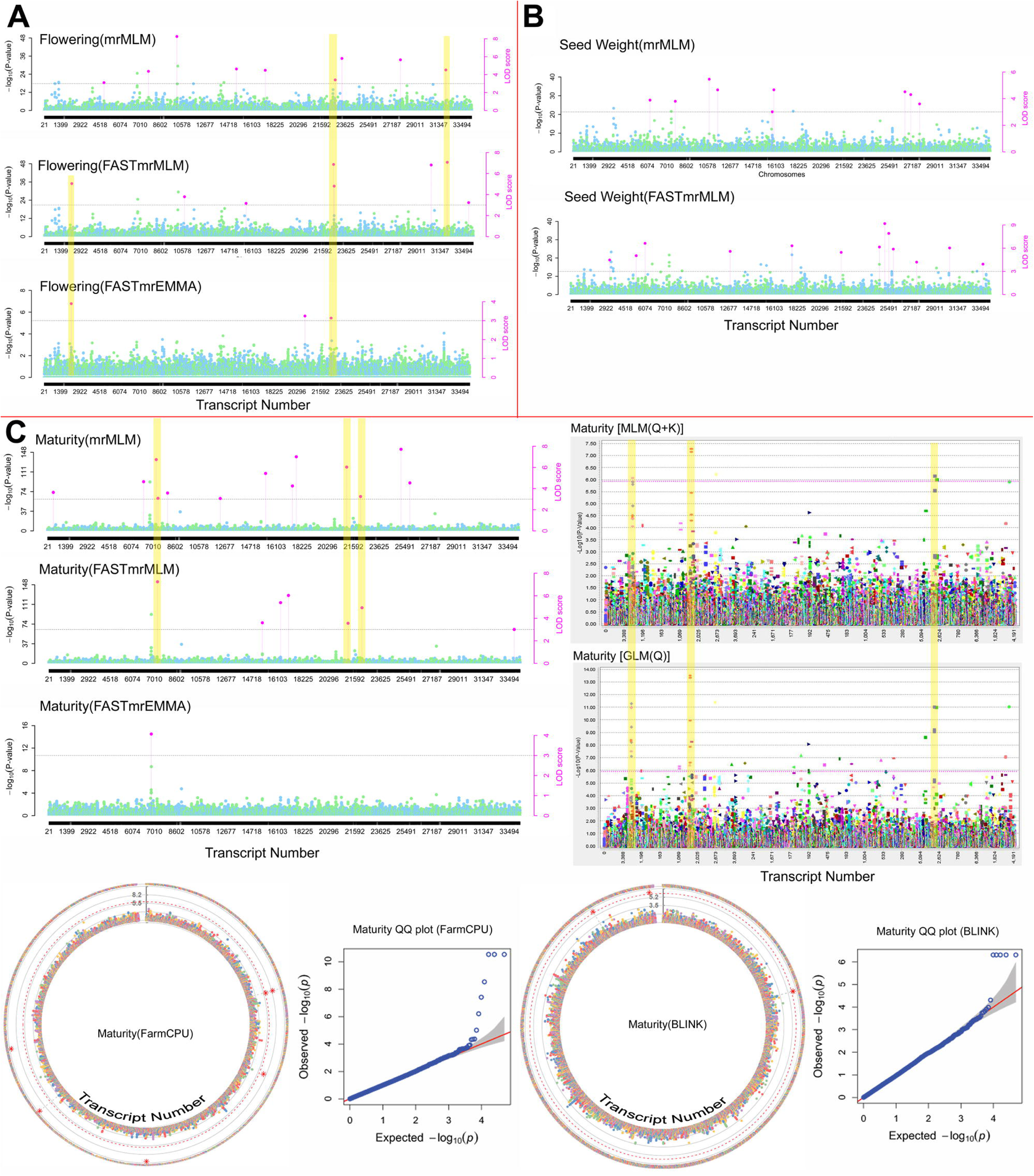
Significantly associated markers for flowering (A), seed weight (B) and maturity (C) shown on the Manhattan plots, predicted by various models using the phenotypic dataset collected from Almora in the year 2020. The highlighted regions show the consistent markers predicted by at least two models. The markers shown on single Manhattan plots for any trait are predicted by ISIS EM-BLASSO for which Manhattan plot is not generated.

**Table 1.**
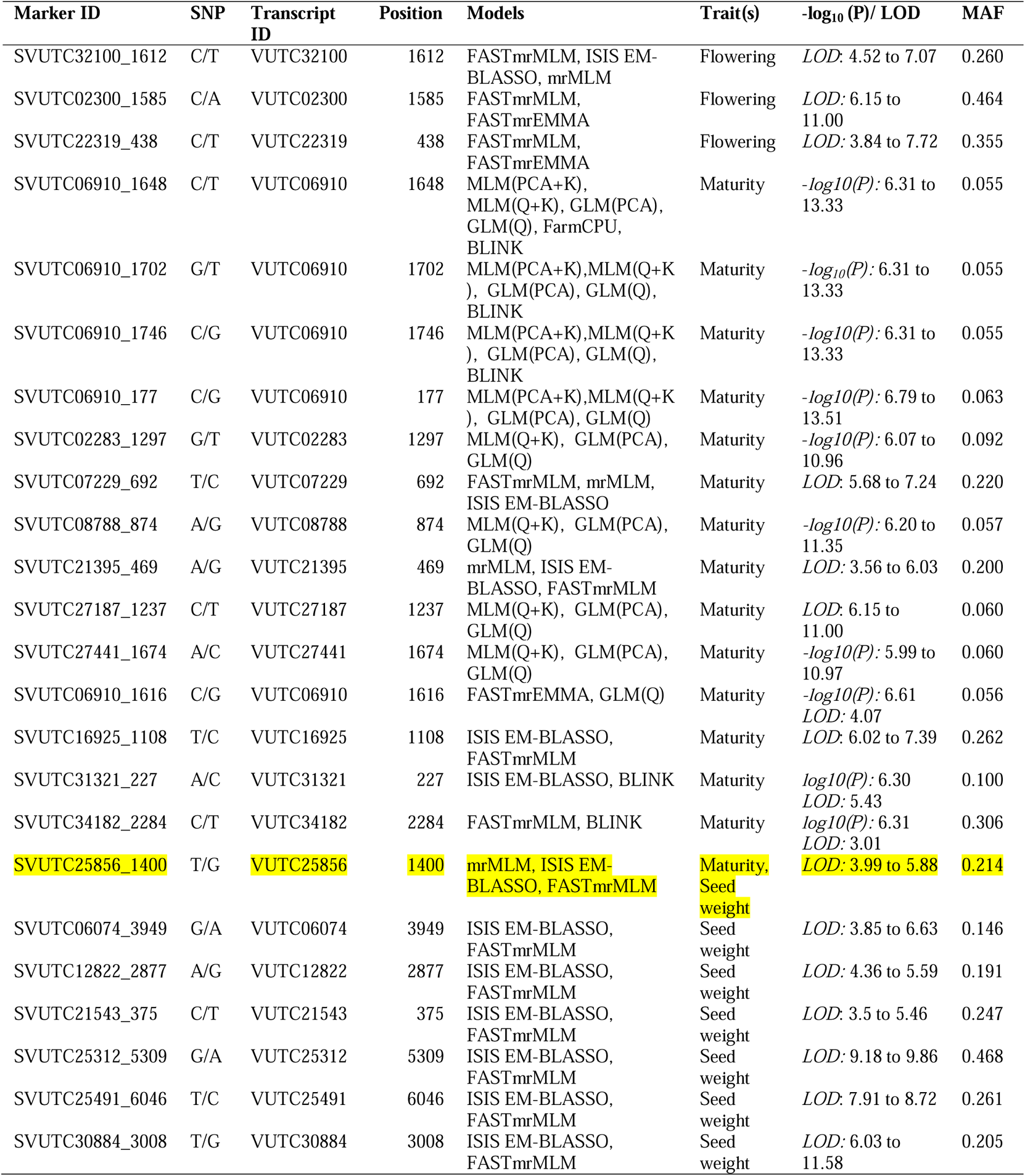
Associated Markers for flowering, maturity and seed weight predicted using phenotypic data of Almora location in the year 2020.

In the case of phenotypic data collected from the Almora location in the year 2021, a total of 11 markers were detected to be associated with different traits (Table 2, Fig. 9). For this dataset, only multi-locus models of mrMLM have detected significant markers which are selected based on LOD score. In this case, 3 markers for flowering, 3 for maturity and 6 for seed weight traits have been predicted where one marker (SVUTC21295_283) was found common for both flowering and maturity traits. Here, all the markers except SVUTC21295_283 were predicted by exactly two different multi-locus models. Though SVUTC21295_283 has been predicted by only one method, it was considered, as it was predicted for two different traits. All the markers were predicted on different transcripts. However, one transcript VUTC22319 was found common in both the years with different marker positions (438 in 2020 and 77 in 2021) and found to be associated with flowering for the datasets of both the years.

**Fig. 9.**
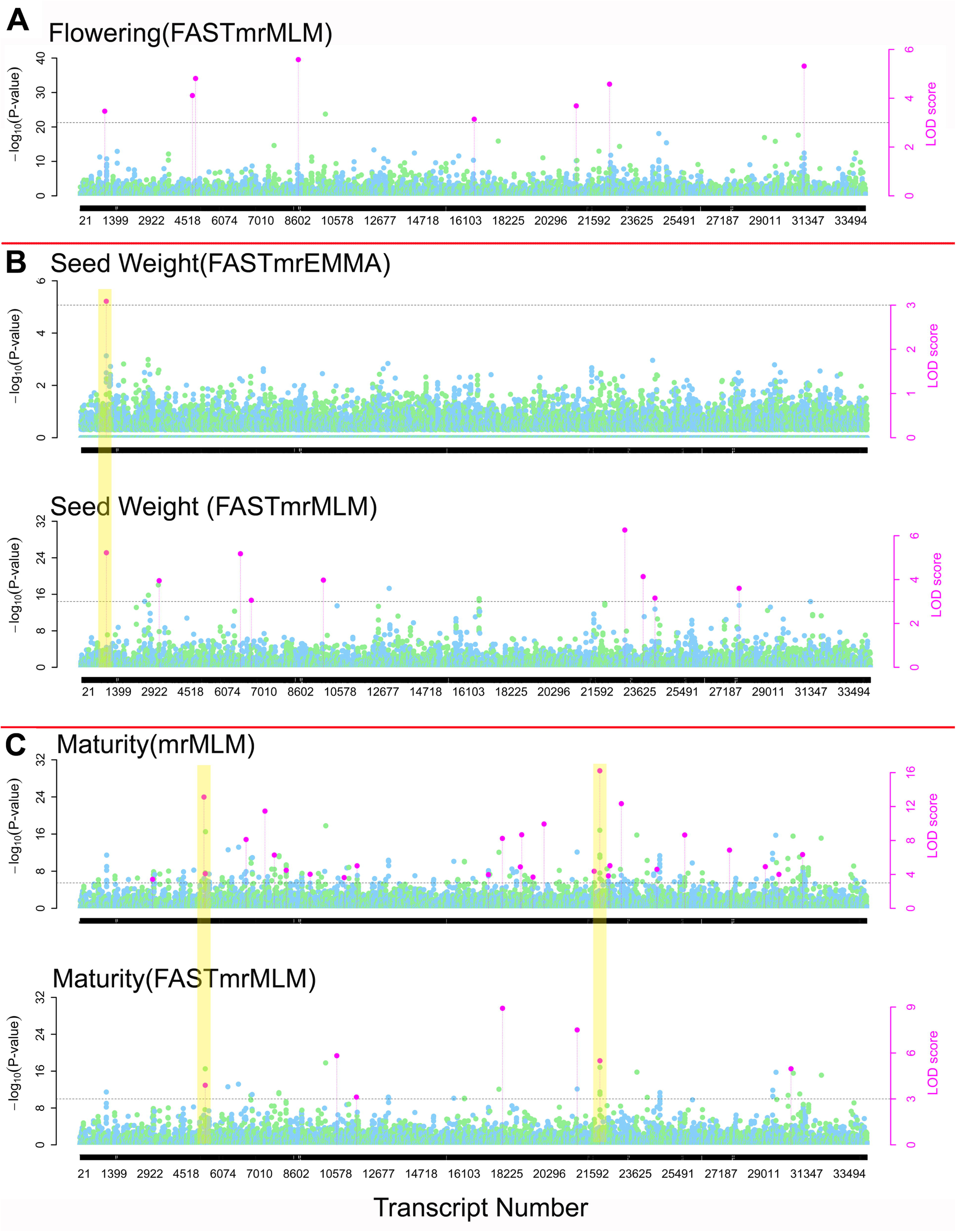
Significantly associated markers for flowering (A), seed weight (B) and maturity(C) shown on the Manhattan plots, predicted by various models using the phenotypic dataset collected from Almora in the year 2021. The highlighted regions show the consistent markers predicted by at least two models. The markers shown on single Manhattan plots for any trait are predicted by ISIS EM-BLASSO for which Manhattan plot is not generated.

**Table 2.** Associated Markers for flowering, maturity and seed weight predicted using phenotypic data of Almora location in the year 2021. *For Almora 2021 only multi-locus models of mrMLM have detected significant markers thus only LOD score is reported.

| Marker ID | SNP | Transcript ID | Position | Models | Trait(s) | LOD* | MAF |
| --- | --- | --- | --- | --- | --- | --- | --- |
| SVUTC21295_283 | G/A | VUTC21295 | 283 | FASTmrMLM | Flowering, Maturity | LOD: 3.69 to 7.50 | 0.110 |
| SVUTC22319_77 | T/C | VUTC22319 | 77 | ISIS EM-BLASSO, FASTmrMLM | Flowering | LOD:4.58 to 5.74 | 0.284 |
| SVUTC31327_3358 | T/G | VUTC31327 | 3358 | ISIS EM-BLASSO, FASTmrMLM | Flowering | LOD:5.32 to 6.18 | 0.165 |
| SVUTC05378_2089 | T/C | VUTC05378 | 2089 | ISIS EM-BLASSO, FASTmrMLM | Maturity | LOD:3.89 to 8.60 | 0.077 |
| SVUTC21831_6083 | G/A | VUTC21831 | 6083 | mrMLM, FASTmrMLM | Maturity | LOD:5.49 to 16.21 | 0.060 |
| SVUTC01132_259 | A/G | VUTC01132 | 259 | FASTmrEMMA, FASTmrMLM | Seed weight | LOD: 3.09 to 5.23 | 0.396 |
| SVUTC06850_2182 | C/A | VUTC06850 | 2182 | ISIS EM-BLASSO, FASTmrMLM | Seed weight | LOD:3.06 to 6.23 | 0.192 |
| SVUTC09951_1960 | C/A | VUTC09951 | 1960 | ISIS EM-BLASSO, FASTmrMLM | Seed weight | LOD:3.97 to 4.78 | 0.3032 |
| SVUTC23058_1028 | A/T | VUTC23058 | 1028 | ISIS EM-BLASSO, FASTmrMLM | Seed weight | LOD:6.26 to 13.15 | 0.140 |
| SVUTC24042_2561 | T/A | VUTC24042 | 2561 | ISIS EM-BLASSO, FASTmrMLM | Seed weight | LOD:4.13 to 5.71 | 0.318 |
| SVUTC24699_1593 | G/T | VUTC24699 | 1593 | ISIS EM-BLASSO, FASTmrMLM | Seed weight | LOD:3.16 to 5.69 | 0.081 |

#### Marker-trait association with Delhi datasets

The marker-trait association analysis for phenotypic data of Delhi in the year 2021 revealed a total of 58 markers out of which majority of the markers(49) were for flowering trait whereas, 4 for maturity and 5 for seed weight were identified (Table 3, Fig. 10). Here, none of the markers were found associated with more than one trait. The 58 markers are located on 24 transcripts, where 7 transcripts contain more than two markers each. The transcript VUTC28154 contains highest (8) number of markers on it. Out of 49 markers of flowering, 43 markers were predicted only by single locus GWAS models. A marker, SVUTC28154_1646 predicted to be associated with flowering has been predicted by 8 out of 10 considered GWAS models.

**Fig. 10.**
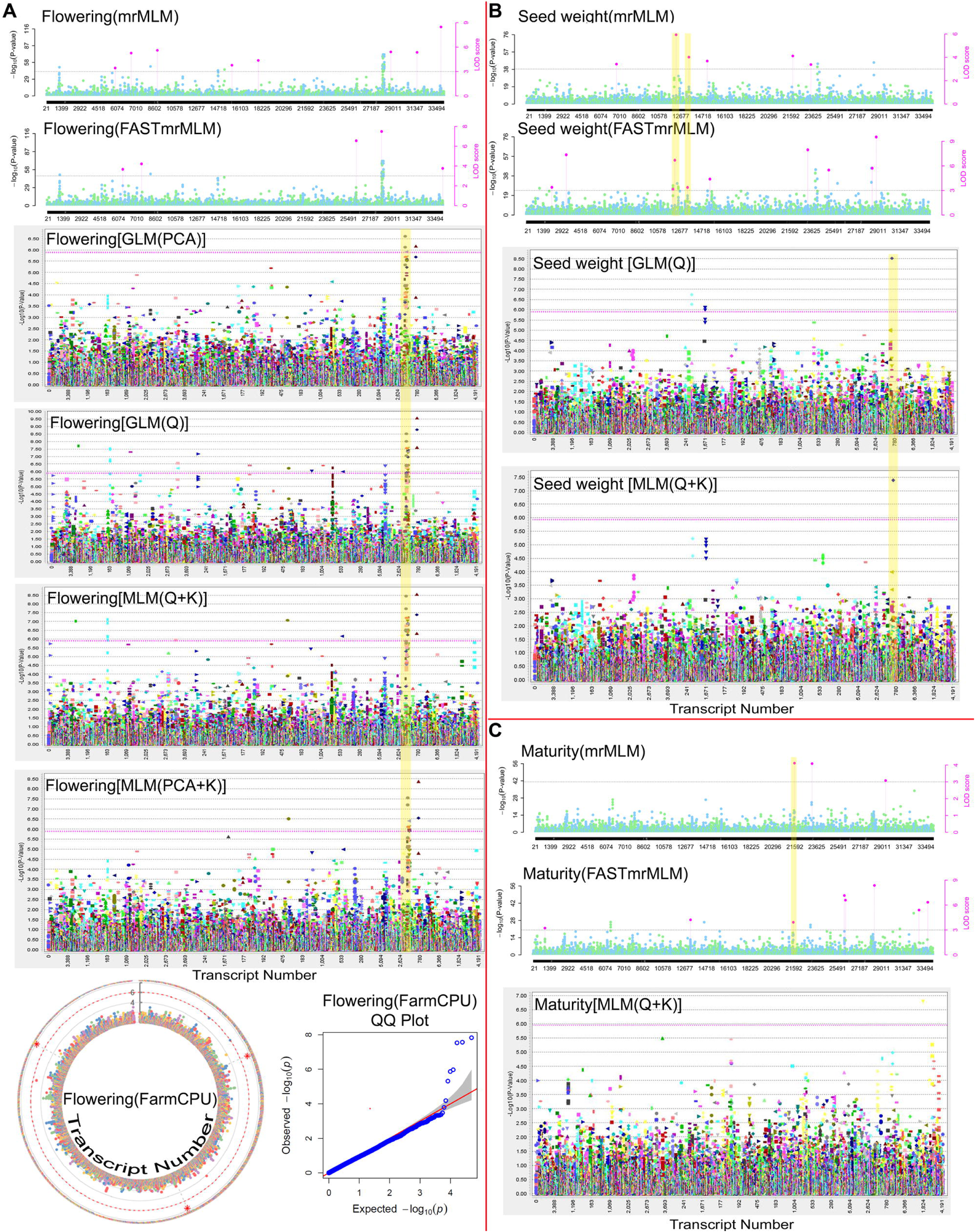
Significantly associated markers for flowering (A), seed weight (B) and maturity(C) shown on the Manhattan plots, predicted by various models using the phenotypic dataset collected from Delhi in the year 2020. The highlighted regions show the consistent markers predicted by at least two models. The markers shown on single Manhattan plots for any trait are predicted by ISIS EM-BLASSO for which Manhattan plot is not generated.

**Table 3.**
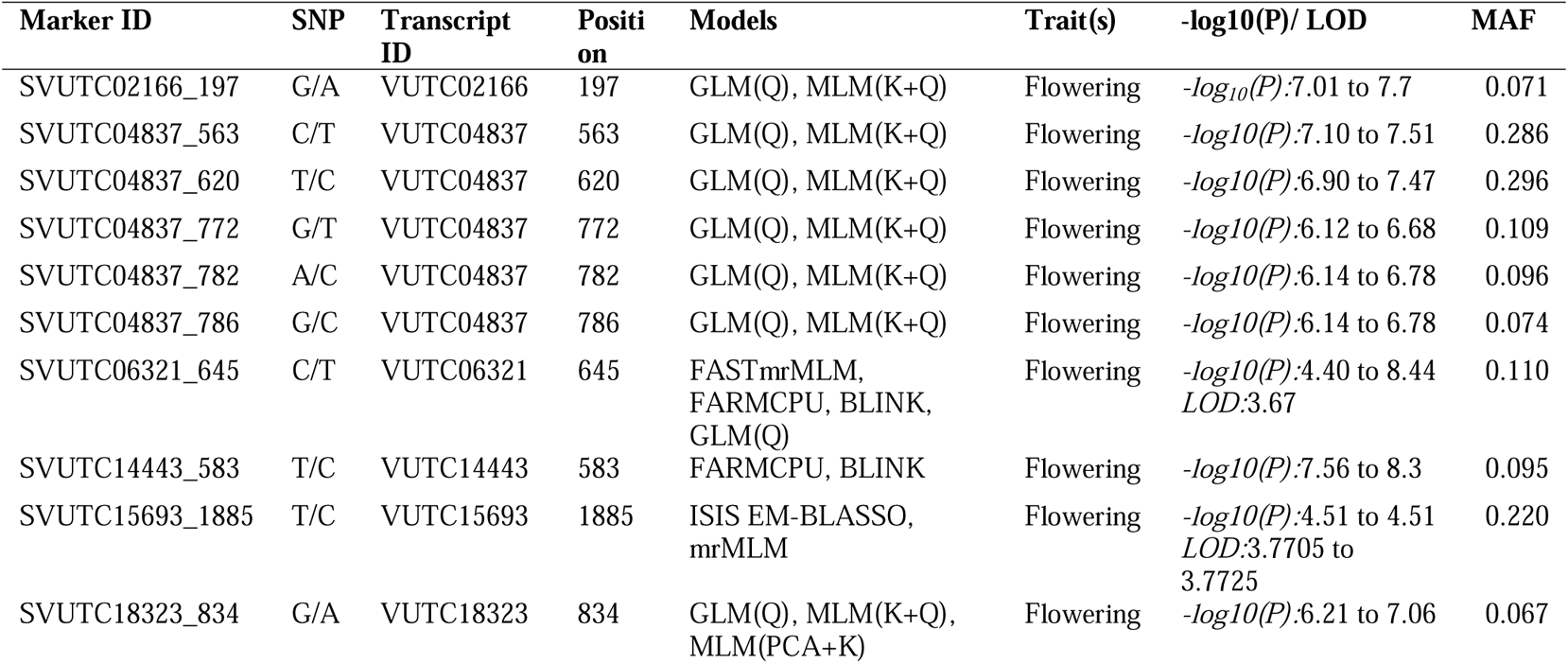
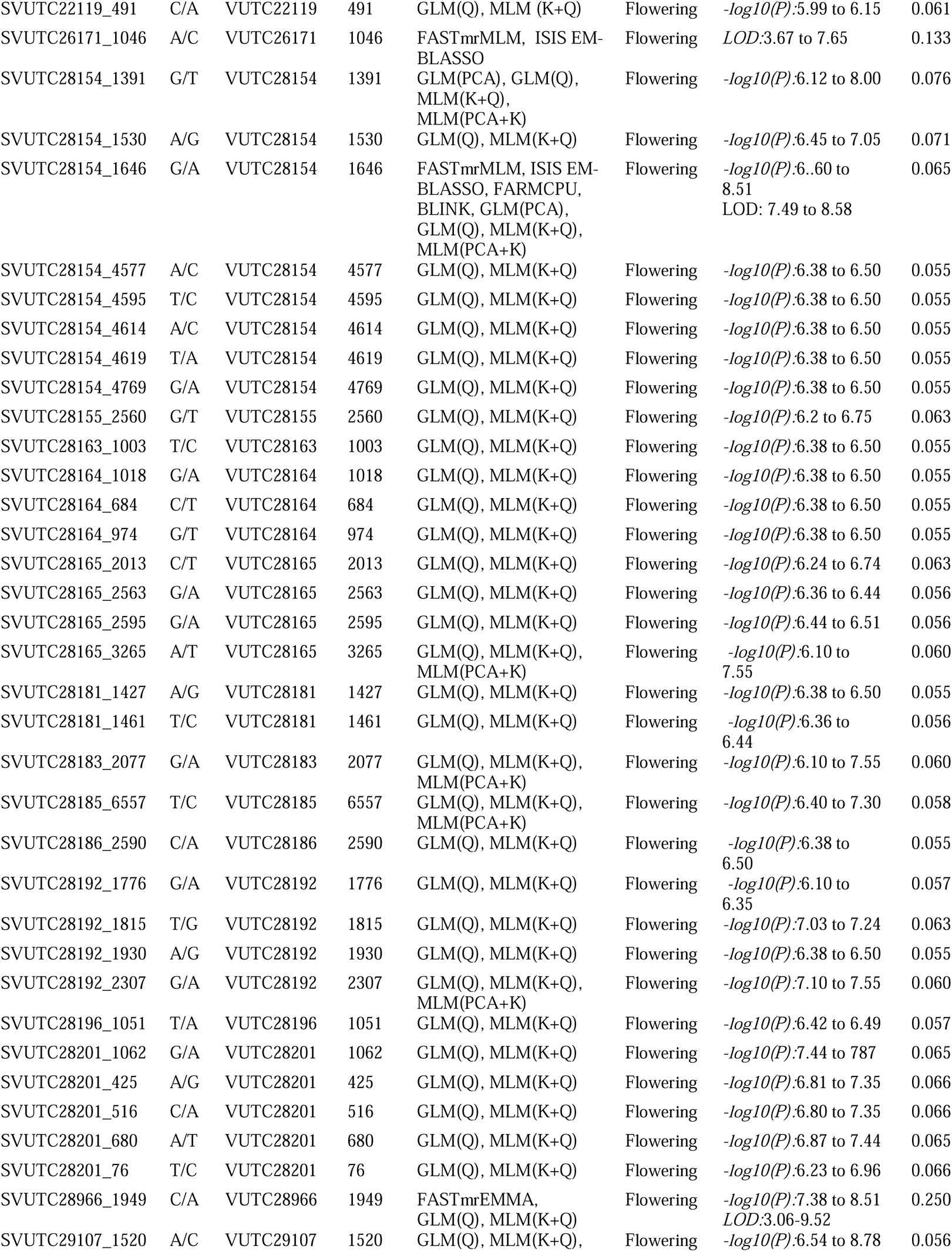
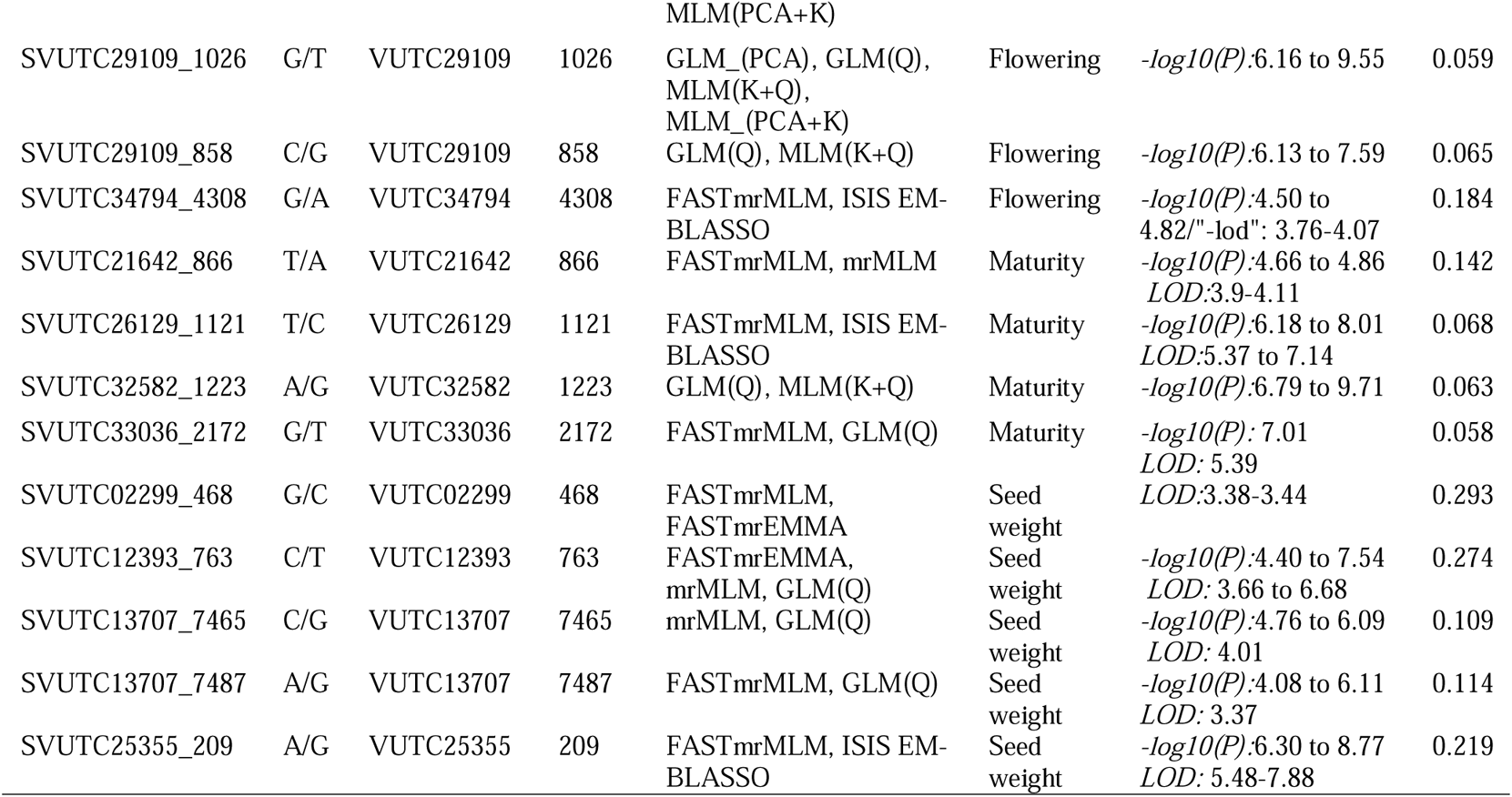
Associated Markers for flowering, maturity and seed weight predicted using phenotypic data of Delhi location in the year 2020.

For the phenotypic data collected from Delhi in the year 2021, the GWAS analysis revealed the association of 49 markers for the three considered traits (Table 4, Fig. 11). Out of these 49 markers, 4 for seed weight, 4 for maturity and 42 were found associated with flowering. Interestingly, one marker (SVUTC25248_8323) being predicted by three methods was found associated with maturity and seed weight. Nine transcripts associated with flowering were identified by at least two markers on them where the transcript VUTC28201 contains highest number of markers on it. A total of 15 markers and 8 transcripts were found common between two years data of Delhi, all of which were observed to be associated with the flowering trait.

**Fig. 11.**
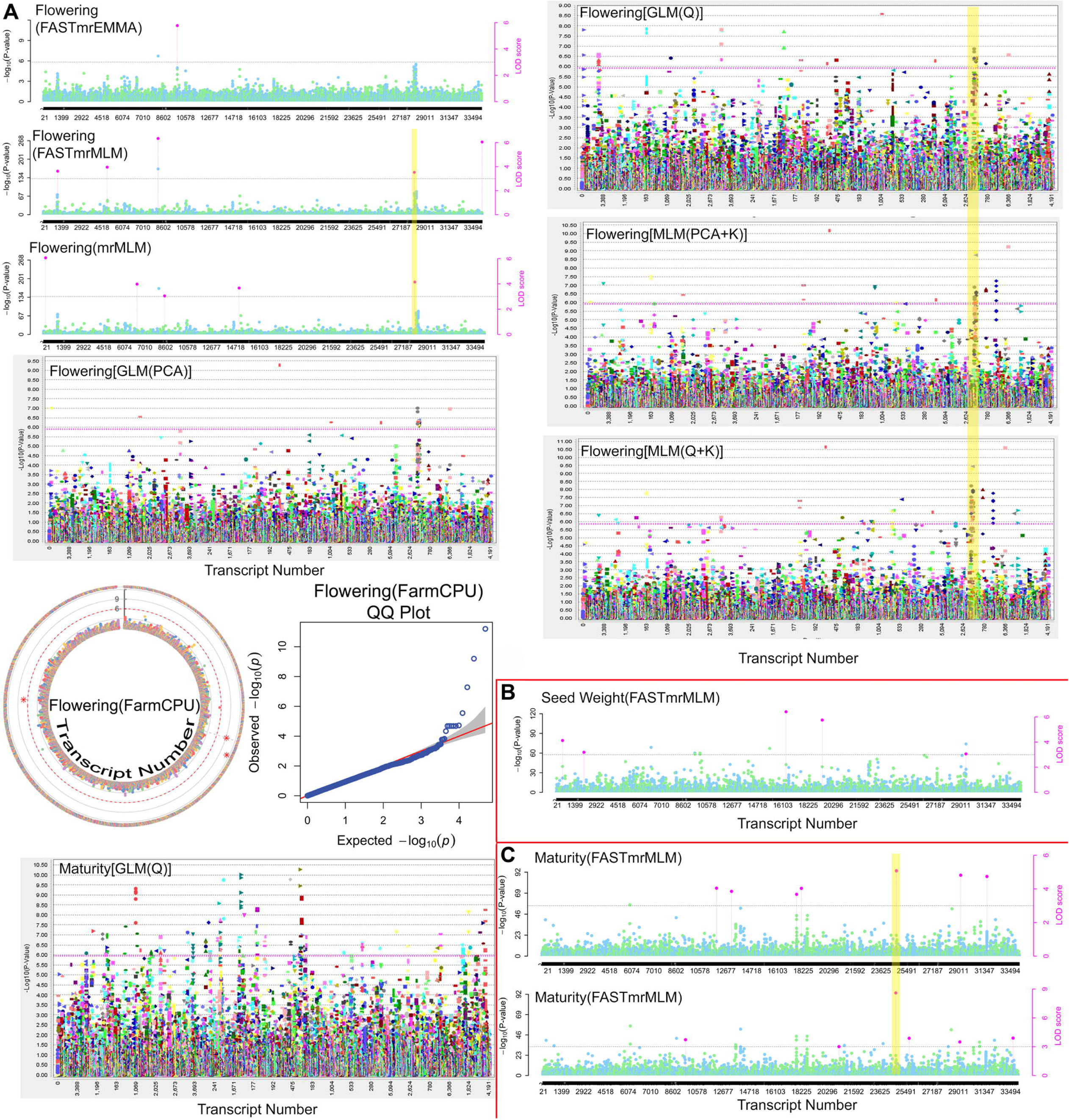
Significantly associated markers for flowering (A), seed weight (B) and maturity(C) shown on the Manhattan plots, predicted by various models using the phenotypic dataset collected from Delhi in the year 2021. The highlighted regions show the consistent markers predicted by at least two models. The markers shown on single Manhattan plots for any trait are predicted by ISIS EM-BLASSO for which Manhattan plot is not generated.

**Table 4.** Associated Markers for flowering, maturity and seed weight predicted using phenotypic data of Delhi location in the year 2021.

| Marker ID | SNP | Transcript ID | Position | Models | Trait(s) | $-\log_{10}(P)$/ LOD | MAF |
| --- | --- | --- | --- | --- | --- | --- | --- |
| SVUTC00230_1049 | A/T | VUTC00230 | 1049 | mrMLM, GLM_PCA,<br>MLM_PCA_K | Flowering | $-\log_{10}(p)$: 6.02 to<br>7.01<br>LOD: 6.063 | 0.113 |
| SVUTC01074_2125 | A/G | VUTC01074 | 2125 | MLM_(PCA+K),<br>MLM(Q+K) | Flowering | $-\log_{10}(p)$: 6.37 to<br>7.09 | 0.118 |
| SVUTC01110_689 | T/C | VUTC01110 | 689 | ISIS EM-BLASSO,<br>GLM(Q) | Flowering | $-\log_{10}(P)$:5.90 to<br>6.31 LOD: 5.10 | 0.070 |
| SVUTC04786_495 | T/A | VUTC04786 | 495 | MLM(PCA+K),<br>MLM(Q+K) | Flowering | $-\log_{10}(P)$:7.37 to<br>7.70 | 0.060 |
| SVUTC04786_499 | A/T | VUTC04786 | 499 | MLM_(PCA+K),<br>MLM_(Q+K) | Flowering | $-\log_{10}(P)$:7.18 to<br>7.52 | 0.060 |
| SVUTC06771_4276 | G/C | VUTC06771 | 4276 | GLM (PCA),<br>MLM(PCA+K) | Flowering | $-\log_{10}(P)$:6.43 to<br>6.55 | 0.153 |
| SVUTC08154_978 | G/A | VUTC08154 | 978 | FASTmrMLM, ISIS EM-BLASSO | Flowering | $-\log_{10}(P)$:7.18 to<br>7.19 LOD: 6.341<br>to 6.3495 | 0.051 |
| SVUTC09869_1881 | C/G | VUTC09869 | 1881 | GLM(Q), MLM(Q+K) | Flowering | $-\log_{10}(P)$:6.28 to<br>7.81 | 0.098 |
| SVUTC09870_149 | C/T | VUTC09870 | 149 | FASTmrEMMA, ISIS<br>EM-BLASSO, FarmCPU,<br>GLM(Q) | Flowering | $-\log_{10}(P)$:6.95 to<br>9.18 LOD: 5.77 to<br>7.12 | 0.092 |
| SVUTC09871_149 | C/T | VUTC09871 | 149 | GLM(Q), MLM(Q+K) | Flowering | $-\log_{10}(P)$:6.11 to<br>6.33 | 0.077 |
| SVUTC15414_2770 | T/C | VUTC15414 | 2770 | GLM(Q),<br>MLM(PCA+K),<br>MLM(Q+K) | Flowering | $-\log_{10}(P)$:6.30 to<br>7.29 | 0.075 |
| SVUTC15414_2854 | A/T | VUTC15414 | 2854 | MLM(PCA+K),<br>MLM(Q+K) | Flowering | $-\log_{10}(P)$:6.16 to<br>6.86 | 0.081 |
| SVUTC15418_711 | C/T | VUTC15418 | 711 | GLM(Q),<br>MLM(PCA+K),<br>MLM(Q+K) | Flowering | $-\log_{10}(P)$:6.30 to<br>7.29 | 0.075 |
| SVUTC17024_838 | C/T | VUTC17024 | 838 | GLM (PCA), GLM(Q),<br>MLM(PCA+K),<br>MLM(Q+K) | Flowering | $-\log_{10}(P)$:6.14 to<br>10.64 | 0.111 |
| SVUTC20322_1284 | A/G | VUTC20322 | 1284 | MLM(PCA+K),<br>MLM(Q+K) | Flowering | $-\log_{10}(P)$:5.99 to<br>6.82 | 0.148 |
| SVUTC20322_1298 | A/T | VUTC20322 | 1298 | MLM(PCA+K),<br>MLM(Q+K) | Flowering | $-\log_{10}(P)$:5.92 to<br>5.99 | 0.160 |
| SVUTC20819_606 | T/C | VUTC20819 | 606 | GLM(PCA), GLM(Q) | Flowering | $-\log_{10}(P)$:6.16 to<br>6.86 | 0.079 |
| SVUTC25024_1324 | A/G | VUTC25024 | 1324 | GLM(PCA), GLM(Q),<br>MLM(PCA+K),<br>MLM(Q+K) | Flowering | $-\log_{10}(P)$:6.23 to<br>6.59 | 0.090 |
| SVUTC25312_4994 | A/G | VUTC25312 | 4994 | ISIS EM-BLASSO,<br>FarmCPU | Flowering | $-\log_{10}(P)$:5.66 to<br>7.27 *LOD*: 4.87 | 0.376 |
| SVUTC28152_1002 | C/T | VUTC28152 | 1002 | GLM(Q), MLM(Q+K) | Flowering | $-\log_{10}(P)$:6.40 to<br>6.49 | 0.065 |
| SVUTC28154_1391 | G/T | VUTC28154 | 1391 | FASTmrMLM,<br>GLM(PCA), GLM(Q),<br>MLM(PCA+K),<br>MLM(Q+K) | Flowering | $-\log_{10}(P)$:4.25 to<br>822 *LOD*: 3.52 | 0.076 |
| SVUTC28154_1530 | A/G | VUTC28154 | 1530 | GLM(PCA), GLM(Q),<br>MLM(PCA+K),<br>MLM(Q+K) | Flowering | $-\log_{10}(P)$:6.17 to<br>7.53 | 0.071 |
| SVUTC28154_1646 | G/A | VUTC28154 | 1646 | GLM(PCA), GLM(Q),<br>MLM(PCA+K),<br>MLM(Q+K) | Flowering | $-\log_{10}(P)$:6.18 to<br>7.19 | 0.065 |
| SVUTC28165_3265 | A/T | VUTC28165 | 3265 | GLM(PCA), GLM(Q),<br>MLM(PCA+K),<br>MLM(Q+K) | Flowering | $-\log_{10}(P)$:6.00 to<br>7.16 | 0.060 |
| SVUTC28183_1588 | T/G | VUTC28183 | 1588 | GLM(PCA), GLM(Q),<br>MLM(PCA+K),<br>MLM(Q+K) | Flowering | $-\log_{10}(P)$:6.00 to<br>9.27 | 0.074 |
| SVUTC28183_2077 | G/A | VUTC28183 | 2077 | GLM(PCA), GLM(Q),<br>MLM(PCA+K),<br>MLM(Q+K) | Flowering | $-\log_{10}(P)$:6.00 to<br>7.16 | 0.060 |
| SVUTC28185_6557 | T/C | VUTC28185 | 6557 | GLM_PCA, GLM_Q, | Flowering | $-\log_{10}(P)$:5.99 to<br>6.38 | 0.058 |
| SVUTC28186_2802 | G/C | VUTC28186 | 2802 | MLM_PCA_K,<br>MLM_Q_K | Flowering | $-\log_{10}(P)$:6.18 to<br>6.26 | 0.071 |
| SVUTC28192_1815 | T/G | VUTC28192 | 1815 | GLM_PCA, GLM_Q,<br>MLM_Q_K | Flowering | $-\log_{10}(P)$:6.04 to<br>7.05 | 0.063 |
| SVUTC28192_2307 | G/A | VUTC28192 | 2307 | GLM(PCA), GLM(Q),<br>MLM(PCA+K),<br>MLM(Q+K) | Flowering | $-\log_{10}(P)$:6.00 to<br>7.16 | 0.060 |
| SVUTC28201_1062 | G/A | VUTC28201 | 1062 | GLM(PCA), GLM(Q),<br>MLM(PCA+K),<br>MLM(Q+K) | Flowering | $-\log_{10}(P)$:6.25 to<br>7.95 | 0.065 |
| SVUTC28201_425 | A/G | VUTC28201 | 425 | GLM(PCA), GLM(Q),<br>MLM(PCA+K),<br>MLM(Q+K) | Flowering | $-\log_{10}(P)$:6.26 to<br>7.90 | 0.066 |
| SVUTC28201_516 | C/A | VUTC28201 | 516 | GLM(PCA), GLM(Q),<br>MLM(PCA+K),<br>MLM(Q+K) | Flowering | $-\log_{10}(P)$:6.17 to<br>7.85 | 0.066 |
| SVUTC28201_680 | A/T | VUTC28201 | 680 | GLM(PCA), GLM(Q),<br>MLM(PCA+K),<br>MLM(Q+K) | Flowering | $-\log_{10}(P)$:6.25 to<br>7.95 | 0.065 |
| SVUTC28201_76 | T/C | VUTC28201 | 76 | GLM(PCA), GLM(Q),<br>MLM(PCA+K),<br>MLM(Q+K) | Flowering | $-\log_{10}(P)$:6.17 to<br>7.82 | 0.066 |
| SVUTC29109_1026 | G/T | VUTC29109 | 1026 | MLM(PCA+K),<br>MLM(Q+K) | Flowering | $-\log_{10}(P)$:6.68 to<br>7.99 | 0.059 |
| SVUTC29109_858 | C/G | VUTC29109 | 858 | MLM(PCA+K),<br>MLM(Q+K) | Flowering | $-\log_{10}(P)$:6.78 to<br>7.50 | 0.065 |
| SVUTC29946_2497 | T/G | VUTC29946 | 2497 | MLM(PCA+K),<br>MLM(Q+K) | Flowering | $-\log_{10}(P)$:6.10 to<br>6.68 | 0.087 |
| SVUTC29946_2507 | A/G | VUTC29946 | 2507 | MLM(PCA+K),<br>MLM(Q+K) | Flowering | $-\log_{10}(P)$:6.64 to<br>7.31 | 0.091 |
| SVUTC29946_2525 | G/C | VUTC29946 | 2525 | MLM(PCA+K),<br>MLM(Q+K) | Flowering | $-\log_{10}(P)$:7.25 to<br>7.25 | 0.090 |
| SVUTC29946_2663 | G/A | VUTC29946 | 2663 | MLM(PCA+K),<br>MLM(Q+K) | Flowering | $-\log_{10}(P)$:6.96 to<br>7.76 | 0.077 |
| SVUTC31155_1020 | A/G | VUTC31155 | 1020 | GLM(PCA), GLM(Q),<br>MLM(PCA+K),<br>MLM(Q+K) | Flowering | $-\log_{10}(P)$:6.23 to<br>10.59 | 0.061 |
| SVUTC12308_931 | T/A | VUTC12308 | 931 | FASTmrMLM, GLM(Q) | Maturity | $-\log_{10}(P)$:4.79 to<br>8.57 LOD: 4.04 | 0.163 |
| SVUTC18444_3244 | C/A | VUTC18444 | 3244 | FASTmrMLM, GLM_Q | Maturity | $-\log_{10}(P)$:4.7 to<br>9.45 LOD: 4.032 | 0.148 |
| SVUTC29349_1348 | G/A | VUTC29349 | 1348 | FASTmrMLM, mrMLM | Maturity | $-\log_{10}(P)$:4.22 to<br>5.60 LOD: 3.49-4.81 | 0.478 |
| SVUTC25248_8323 | G/A | VUTC25248 | 8323 | FASTmrMLM, mrMLM,<br>ISIS EM-BLASSO | Maturity,<br>Seed<br>weight | $-\log_{10}(P)$:5.22 to<br>9.50<br>LOD: 4.44-8.59 | 0.159 |
| SVUTC00580_1033 | T/C | VUTC00580 | 1033 | FASTmrMLM, ISIS EM-BLASSO | Seed<br>weight | $-\log_{10}(P)$:3.75 to<br>4.88 LOD: 3.05 to<br>4.12 | 0.085 |
| SVUTC16440_582 | A/G | VUTC16440 | 582 | FASTmrMLM, ISIS EM-BLASSO | Seed<br>weight | $-\log_{10}(P)$:4.59 to<br>7.25 LOD: 3.84 to<br>7.25 | 0.074 |
| SVUTC19671_1032 | G/A | VUTC19671 | 1032 | FASTmrMLM, ISIS EM-BLASSO | Seed<br>weight | $-\log_{10}(P)$:4.48 to<br>6.57 LOD: 3.74 to<br>5.75 | 0.087 |

### Associated markers and corresponding transcripts

From the overall analysis, 87 transcripts were found associated with the traits having 127 markers in total from all the datasets considered. From the datasets of two locations in two consecutive years, neither any marker nor any transcript could be found in common for all. However, 15 markers were noticed to be common between Delhi-2020 & Delhi-2021 datasets (Fig. 12A). Transcripts are concerned, one transcript (VUTC22319: flowering) between Almora-2020 & Almora-2021 datasets and 8 transcripts (VUTC28154, VUTC28165, VUTC28183, VUTC28185, VUTC28192, VUTC28201, VUTC29109, VUTC28186; all associated with flowering trait) between Delhi-2020 & Delhi-2021 datasets were identified to be common (Fig. 12B). Interestingly, one transcript (VUTC25312: flowering) was also found common between Almora-2020 and Delhi-2021 datasets, even though, no marker was identified to be common between these two datasets.

**Fig. 12.**
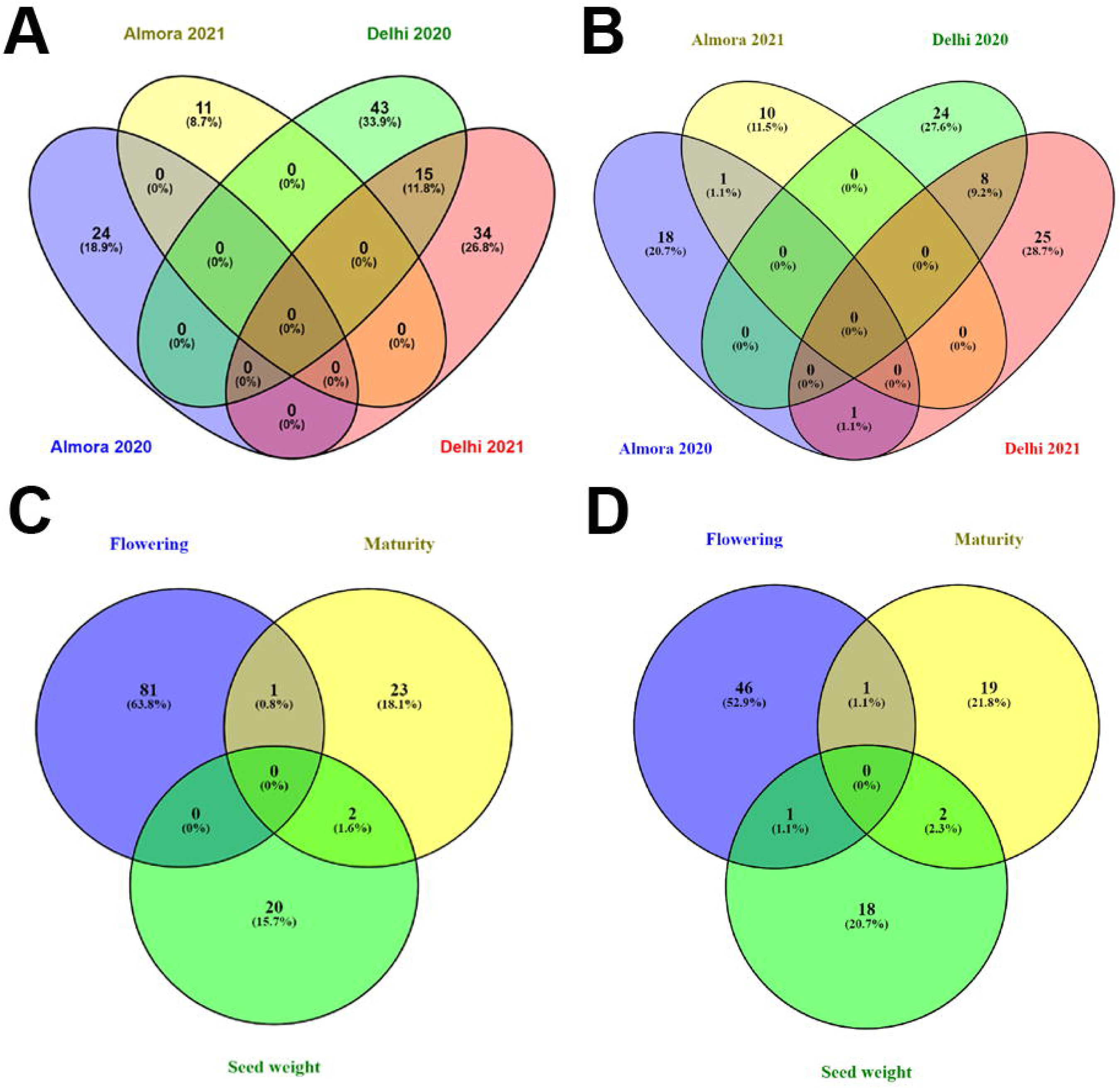
Common markers and transcripts identified for different traits and different datasets. (A) Common markers for datasets, (B) common transcripts for datasets (C) common markers for traits (D) common transcripts for traits.

From 3 out of 127 markers and 4 out of 87 transcripts were identified to be associated with more than one trait. One marker (SVUTC21295_283) between flowering and maturity and 2 markers (SVUTC25856_1400, SVUTC25248_8323) between flowering and seed weight were identified (Fig. 12C). Further, one transcript (VUTC21295) between flowering and maturity, 2 transcripts (VUTC25856, VUTC25248) between maturity and seed weight and one transcript between flowering and seed weight (VUTC25312) have been identified (Fig. 12D). However, later one was predicted for two different datasets (Almora-2020 and Delhi-2021 with two different markers *i.e.,* SVUTC25312_5309 and SVUTC25312_4994, respectively). These common markers and transcripts are expected to help understand the interrelation between the considered traits in terms of the governing genes.

### Annotation of associated transcripts

The annotation of transcripts in terms of chromosomal localization (Table S3) and molecular function (Table S4) has revealed many functional proteins related with the associated traits. From the BLAST results, for majority of transcripts, top hits were identified against the proteins of *V. umbellata* and *Vigna anguilaris.* BLAST2GO revealed GO terms for each transcript sequence. The major GO terms for biological processes obtained as annotations for 87 transcripts are presented in the form of pi-chart in the Fig. 13. A hierarchical representation of GO terms for all the associated transcripts is given in the Fig. S1.

**Fig. 13.**
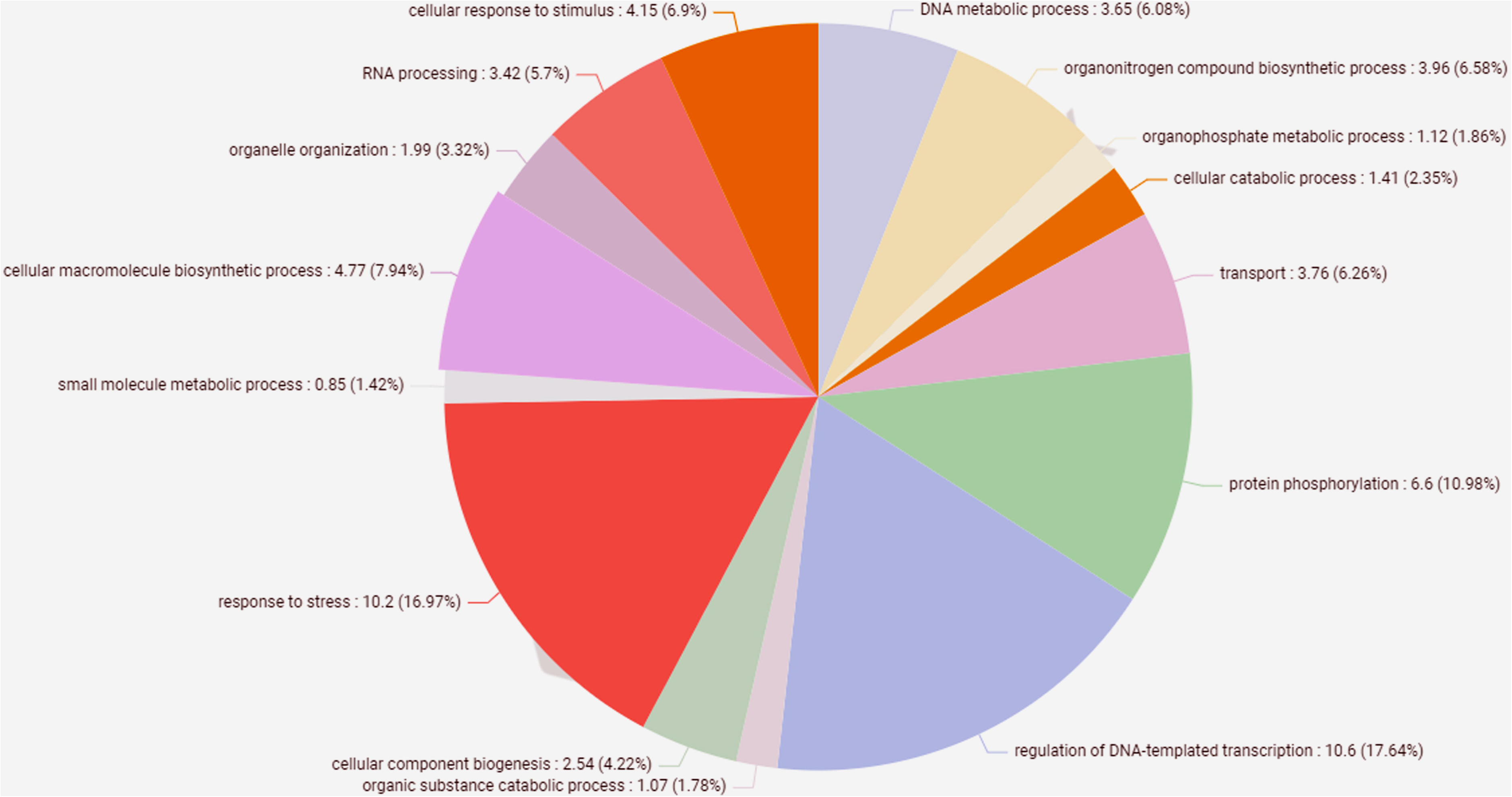
The major gene ontology terms for biological processes obtained as annotations for 87 transcripts presented in the form of pi-chart.

### Chromosomal localization of markers

The exact chromosomal positions of the associated markers were revealed from the ungapped alignment between marker sequences (101 bp; SNP at 51^st^ position) and chromosomal sequences of the recently released rice bean cultivar FF25. The trait-wise associated markers mapped on to different chromosomes of rice bean is shown in Fig. 14. All the associated markers were found distributed over all the chromosomes of rice bean. Most of the markers associated with flowering trait are found on chromosome 1. The highest number of markers for maturity was found on chromosome 11 whereas highest number of markers for seed weight was found on chromosome 5. On chromosome 9, only two markers for flowering were identified. A set of 35 markers were identified on the chromosome 1 with in a distance of 80.83 Kb at 41.6 Mbp to 42.4 Mbp.

**Fig. 14.**
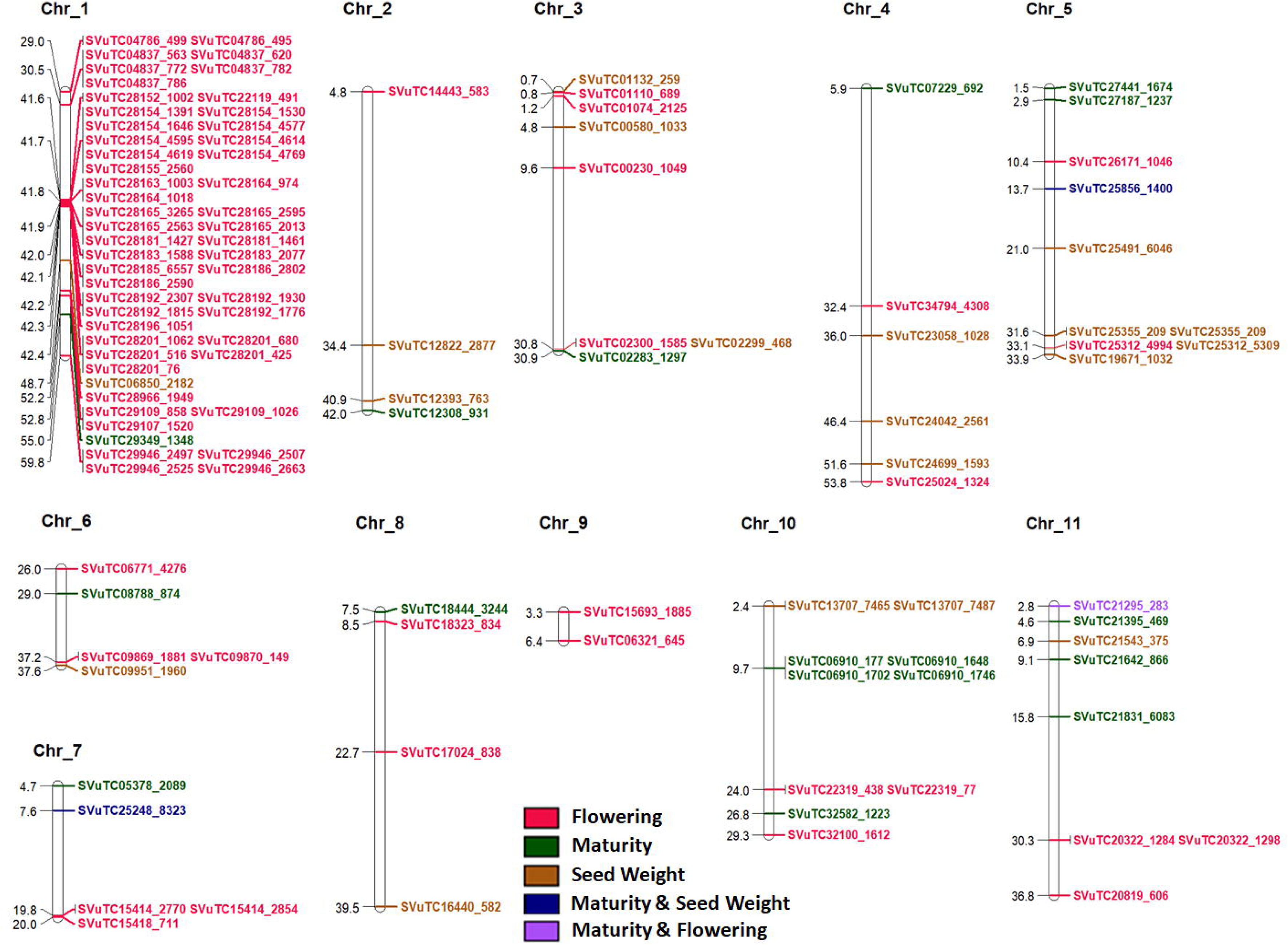
The trait-wise associated markers mapped on to different chromosomes of the rice bean cultivar FF25.

### Inter-species synteny based on chromosomal localization of associated transcripts

The associated transcripts when mapped on to the recently released genome of rice bean (Guan *et al*., 2022); with >= 95% query coverage all the associated transcripts were mapped on to the chromosomes (Table S3). 81 out of 87 transcripts were identified with 100% percentage identity and >=96% query coverage. Further, the mapping of the 87 transcripts on to the chromosomes of different *Vigna* species revealed their inter-species chromosomal locations which assisted in establishing a synteny between all the considered *Vigna* genomes in terms of the trait-associated transcripts (Fig. 15). The pairwise synteny between *V. umbellata* and other *Vigna* species revealed the inter-species chromosomal syntenic relationship. Based on these pair-wise relationships with *V. umbellata,* the synteny between other *Vigna* species was also established (Fig. 15).

**Fig. 15.**
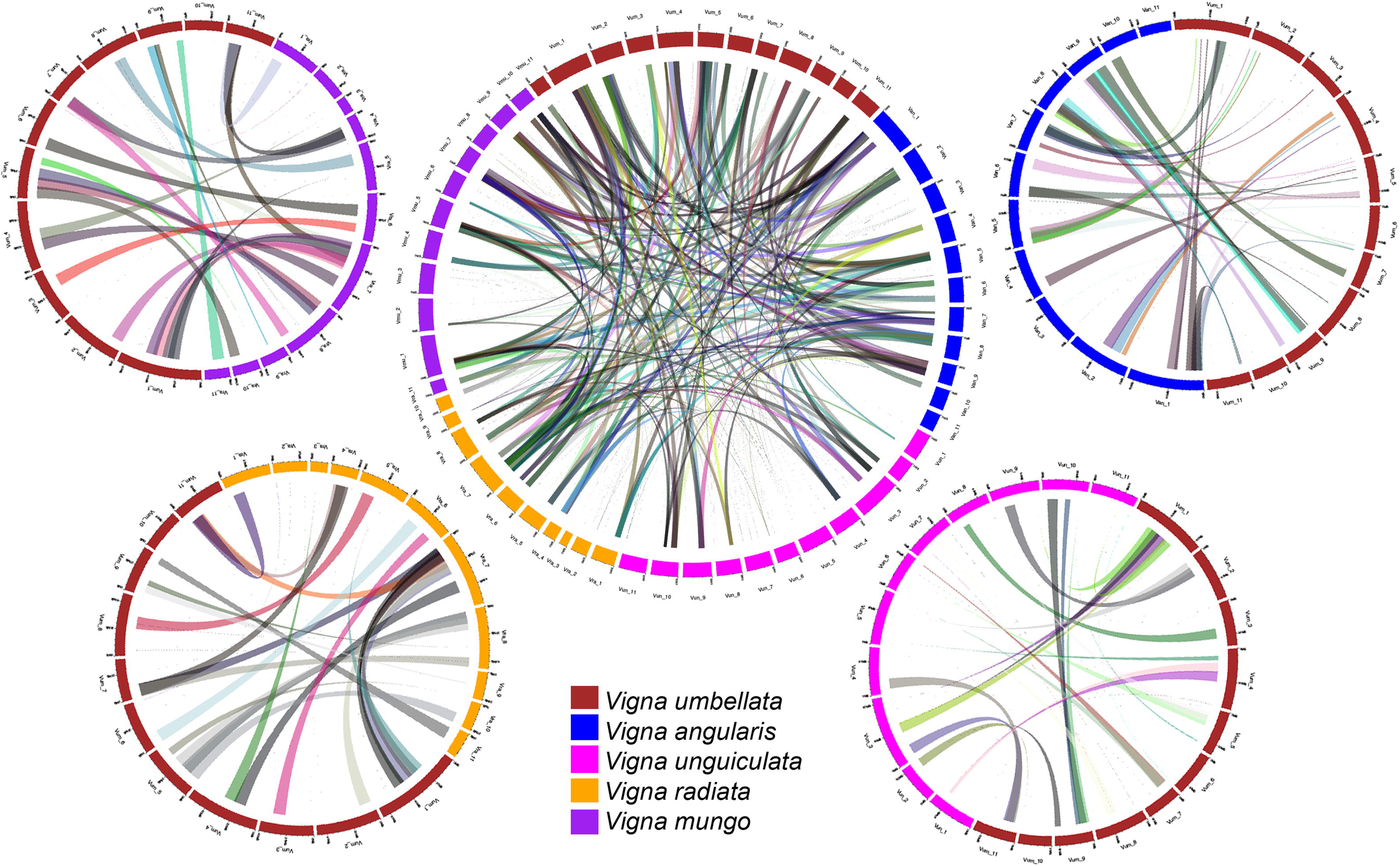
Synteny between all the considered *Vigna* genomes in terms of the trait-associated transcripts along with pair-wise synteny between *Vigna umbellate* and other *Vigna* species.

### Expression analysis of associated transcripts

The expression pattern of genes represented by 87 transcripts for the stages like inflorescence, 5 days post anthesis and 10 days post anthesis is given in the Fig. 16. Most of the associated transcripts for flowering were found enriched with the reads from sample taken at inflorescence and 5 days post anthesis stages. However, the few transcripts associated with maturity and seed weight were also observed to be enriched with the reads from all three stages suggesting their expression during the entire process of flowering to maturity.

**Fig. 16.**
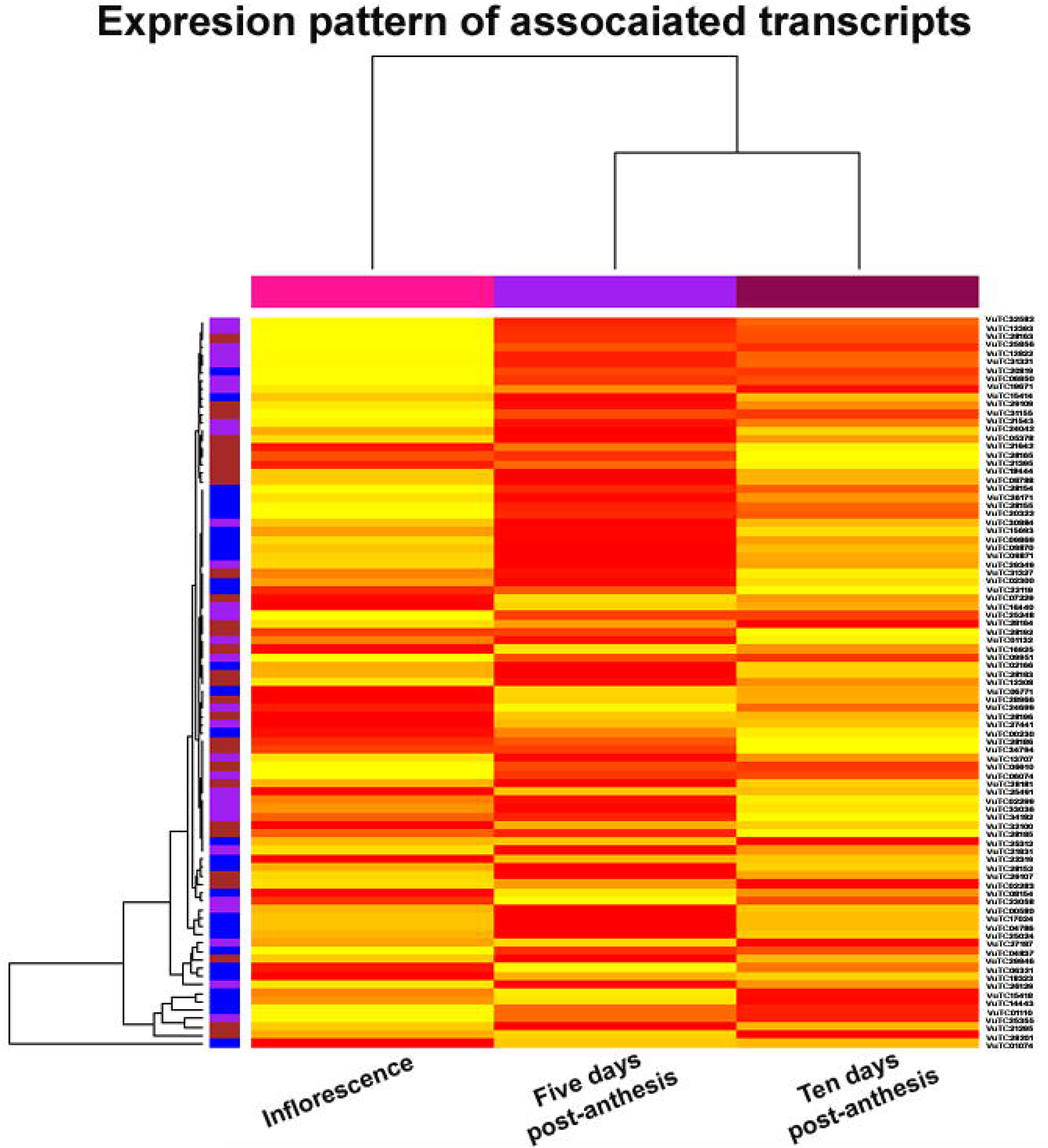
The expression pattern of genes represented by 87 transcripts for the stages like inflorescence, 5 days post anthesis and 10 days post anthesis.

## Discussion

Rice bean being an underutilized crop with high nutritional value bears the potential to contribute to the food and nutritional security across the globe. Despite of these merits, it has been a neglected crop worldwide and remained underutilized. It has received minor attention from breeders and researchers and thus, a little research carried out on this crop. Due to negligence, it also suffers from wild and disadvantageous traits (Pattanayak *et al*., 2019). However, now its importance is being understood and several research projects are undertaken for the genetic improvement of this crop. The genomes released by Chinese Academy of Agricultural Sciences, Beijing, China (Guan *et al*., 2022) at chromosomal level and International Centre for Genetic Engineering and Biotechnology, New Delhi, India at scaffold level (Kaul *et al*., 2022) have increased the scope of genomic research on rice bean.

Information on population structure in GWAS analysis is often incorporated to avoid false marker-trait associations. Q-matrix generated through STRUCTURE and PC scores from PCA help inferring the population structure from the genotypic data in GWAS analysis (Abraham and Inouye, 2014). STRUCTURE and PCA indicated the classification of genotypes into three subpopulations (Population 1-3) that has also corroborated with the genotypic cluster derived through TASSEL (Fig. 6). Population 1 contains 53 genotypes with 80 days on an average to 50% flowering, 123 days on an average to 80% maturity and an average 100-seed-weight of 6.96 gm. Further, the Population 2 was noticed to have 32 genotypes with average values of 74 days, 116 days, 6.61 gm for 50% flowering, 80% maturity and 100 seed weight respectively. Furthermore, the population 3 was observed to have a group of 15 genotypes with average values of 69 days, 106 days and 6.54gm for 50% flowering, 80% maturity and 100-seed-weight respectively. This implies the population 1 contains mostly the late flowering and late maturity genotypes whereas population 2 and 3 contain most of the genotypes of early and normal flowering type.

The phenotypic data from two locations in two consecutive years when subjected to ANNOVA, data for most of the traits were found significantly different (Fig. 2). Seed weight and maturity data for Almora 2020 and 2021 were not found significantly different. Further, the maturity data for Delhi 2020 and Delhi 2021 were also not observed to be significantly different. Interestingly, the flowering data for Almora 2020 and Delhi 2021 were also found to differ insignificantly. The significant difference in the phenotypic data is expected for different locations as Almora has a hilly topography and received high rain, whereas Delhi has a plain topography and had much less rain during the crop season. However, the difference in the data during consecutive years in the same location could be due to unpredictable climate change at these two locations in consecutive years.

In cross pollinated crops LD decays at short distance as compared to self pollinated crops (Dinesh *et al*., 2016, Yu *et al*., 2014). Rice bean is a highly cross pollinated crop, thus, we have also observed decay of LD at short distances. The LD decay at an *r*^2^ cutoff of 0. 2 was observed to be 1.5 Kb (Fig. 7) suggesting a high genetic diversity in the genotypes taken into consideration. Further, the LD decay at short distances can also be expected with the transcript data as mature mRNA lacks introns. Thus, markers identified through GWAS with in a distance of 1.5 kb are expected to be associated with similar or related traits and are expected to be inherited together with a little chance of contemporary recombination. Though, several GWAS models exist with their own advantages and disadvantages, all models cannot well fit with all datasets because of the different distribution patterns in the datasets. We have implemented here 10 different models and selected markers on the basis of their threshold score. The markers were further screened if they are predicted by at least two models. Our approach was to minimize the false positive as well as not to discard biologically important markers even if they are not predicted by high number of methods.

In spite of applying several models we could not find any marker for any trait found consistently with all dataset. However, we observed few markers and corresponding transcripts consistent with two datasets (Fig. 12). As rice bean is a highly cross pollinated crop, the associated genes may very location to location depending upon the difference in the topography, latitude, and longitude and crop duration. This assertion is supported by the findings of Guan et al., 2022 that reported distinct association signals for the different locations. They conducted GWAS analysis for various traits from which they have also included two of the traits *i.e.,* flowering and seed weight that was considered in our study. Two of their reported markers, Chr6:25.86Mb and Chr10: 28.15Mb have shown association with flowering. We have also observed the association two markers (SVUTC06771_4276: Chr6:26.03Mb and SVUTC32100_1612:Chr10:29.31 respectively) for flowering within the vicinity (1.5Mb) of their reported markers. Additionally, we found four markers (SVUTC06850_2182: Chr1:48.73Mb, SVUTC25856_1400: Chr5:13:66Mb, SVUTC09951_1960:Chr6:37.65Mb, SVUTC25248_8323:Chr7:7.58Mb) for seed weight nearby the four markers (Chr1:48.26, Chr5:14.78, Chr6:36.35, Chr7: 6.14, respectively) reported by Guan *et al*. (2022) for seed weight. However, two of these markers (SVUTC25856_1400:Chr5:13.66Mb, SVUTC25248_8323: Chr7:7.58Mb) identified in our study was also found to be associated with maturity trait. Isemura *et al*., 2010 have also reported markers for seed weight on the linkage groups considering inter specific populations, however, our results could not be corroborated with their findings as exact correlation between the linkage groups and chromosomes is unknown. In the chromosomal localization of the genes we found most of the flowering genes on the chromosome 1(Fig. 14) and the role of chromosome 1 in controlling the flowering time has been reported in common bean (Gu et al., 1998; Kwak et al., 2008; González et al., 2016). It is worth mentioning here that chromosome 1 of cow pea corresponds to the chromosome 1 of common bean (Lonardi et al., 2019) and chromosome 1 of rice bean (Guan et al., 2022), which indicates the correspondence between common bean and rice bean genome in terms of chromosome 1. The inter species syntenic relationship (Fig. 15) based on associated transcripts revealed the higher closeness of rice bean with *V. angularis*, *V. mungo* and *V. radiata* than *V. unguiculata*. The same has also been depicted in Guan et al., 2022 and Pattanayak *et al*., 2019.

With the datasets collected from of Almora SVUTC25856_1400 was found to be associated with maturity and seed weight traits. The corresponding transcript (VUTC25856) containing the above mentioned SNP has shown its similarity with transcription factors bHLH149, bHLH147 and PIF3. The GO annotation of this transcript revealed its involvement in the biological processes like regulation of DNA-templated transcription, response to stress, response to oxygen-containing compound with two functions: transcription cis-regulatory region binding and protein dimerization activity. The Basic Helix-Loop-Helix (bHLH) transcription factors regulate the expression of the genes having a broad range of functions in biosynthesis, metabolism and transduction of plant hormones (Hao *et al*., 2021). The phytochrome-interacting factor (PIF3) belongs to bHLH family of proteins that has a specific function in light-induced developmental processes (Ni *et al*., 1998). During seed maturation in light conditions, the amount of abscisic acid in seeds and sensitivity of seeds to abscisic acid is influenced (Contreras *et al*., 2008) that has an important role in the biosynthesis of storage compounds in the embryo, seed dormancy, and the inhibition of precocious germination (McCarty, 1995; Finkelstein et al., 2002; Kanno et al., 2010). Besides, Cheng *et al*. (2014) reported that mutants of *Arabidopsis* deficient with abscisic acid produced seeds with increased size, mass, and embryo cell number. Thus, transcript (VUTC25856) similar to PIF3 with bHLH domain is expected to regulate the abscisic acid dependent behavior of seeds during light-induced developmental processes. Another marker, SVUTC06910_1648 was found for maturity and predicted by 6 out of 10 methods. However, significant annotation for this transcript could not be found as it showed similarity with a predicted protein of *V. umbellata*(XP_047172852) with uncharacterized function. The protein superfamily was identified to be *pepsin_retropepsin_like* having members with aspartic protease function. In *A. thaliana* anther, over expression of such an aspartic protease belonging to this superfamily resulted inhibition of dehiscence process (Marciniak and Przedniczek, 2021). Thus, the transcript VUTC06910 found uncharacterized can be characterized as a member of pepsin_retropepsin_like superfamily having aspartic proteases activity and inhibitory function in pod dehiscence process during maturity in rice bean.

A marker SVUTC21295_283 annotated to have aldo-keto reductase function was found to be associated with maturity and flowering. The aldo-keto reductase enzymes reduce carbonyl substrates like sugar aldehydes, keto-steroids and keto-prostaglandins (Penning, 2015). The D-GalUA reductase, an aldo-keto reductase, plays an important role in the conversion of D-galacturonic acid to L-galactonate which is finally converted to L-ascorbate (in two subsequent steps) in the D-Galacturonic acid pathway for ascorbate biosynthesis in plants (Ishikawa *et al*., 2018). Kotchoni *et al*., 2009 reported that the alterations in the endogenous ascorbic acid content affect flowering time in *Arabidopsis* and they noticed that the artificial increase in ascorbic acid delayed the flowering. Delaying in flowering also affects the days to maturity. Further, vitamin C/ascorbic acid content of rice bean is 15.33–28.23 mg (per 100 g) being second highest in major pulse crops (Tripathi *et al*., 2021) next to pigeon pea (43 mg/100 g). Thus, the transcript VUTC21295 having an aldo-keto reductase function might be playing a regulatory role in ascorbate biosynthesis in rice bean having an effect on its flowering and maturity. Another transcript VUTC22319 was identified with two markers at the positions 438 and 77 with 2020 and 2021 datasets respectively for flowering. The transcript showed similarity with STS14 and pathogenesis-related proteins. The STS14-like proteins have been reported to be highly expressed in the pistil of potato having pathogenesis related function, possibly, for the protection or guidance of the pollen tubes through the pistil (Van Eldik *et al*., 1996).This suggests transcript VUTC22319 might have a similar kind of role in rice bean.

Delhi dataset is concerned, 8 markers were identified on the transcript VUTC28154 and SVUTC28154_1646 was predicted by 8 out of 10 models. The transcript showed similarity with putative phospholipid-transporting ATPase 9 and its GO annotation revealed its involvement in the phospholipid translocation process. Zhou *et al*. (2016) reported the up-regulation of *phospholipid-transporting ATPase* 9 and inferred that *phospholipid-transporting ATPase* 9 along with other phospholibpases may play a part in accelerating the pollen tube aging in *Pyrus bretschneideri*. With this dataset another transcript VUTC25248 was found to be associated with maturity and seed weight traits. The transcript showed similarity with a protein, retrovirus-related Pol polyprotein from transposon TNT 1-94 which is involved in the biological processes like DNA integration and RNA phosphodiester bond hydrolysis with the molecular functions of RNA binding and ribonuclease activity. Cappetta *et al*., (2021) et al., reported that the Retrovirus-related Pol polyprotein from transposon TNT1-94 is associated with heat stress in tomato. Besides, the gene has also been reported to be upregulated in lentil during heat stress (Singh *et al*., 2019). Yamamoto *et al*. (2008) reported that Heat stress (28 to 30°C) has the potential to accelerate plant maturity by reducing seed germination and maturity periods. In the present invesigation, rice bean was grown during kharif season in Delhi and during this period the crop was exposed to heat stress due to hot climate of Delhi. Thus, the Retrovirus-related Pol polyprotein from transposon TNT 1-94 might be playing a role in response to heat stress induced maturity in rice bean. Five markers were identified on VUTC28201 for flowering. The transcript annotation shows its similarity with pectin acetylesterase 8-like protein having role in cell wall organization and pectin acetylesterase activity. However, expression of pectin acetylesterase 8 of *Arabodopsis thaliana* has been reported to be highly regulated during plant growth and development and expressed in different flowering parts and stages(pollen, inflorescence meristem, stamen, petal differentiation and expansion stage) suggesting a role in the control of the degree of acetylation of pectins (Philippe *et al*., 2017, Schmid *et al*., 2005).

Among the top 10 expressed transcripts in all the three stages (inflorescence, 5 days post anthesis and 10 days post anthesis), 8 transcripts are found common. Out of these 8 transcripts, associations of 5 transcripts with flowering, one with both flowering and maturity, one with maturity, and one with seed weight have been revealed. The transcripts associated with only flowering were found to encode heat shock cognate protein 80(HSC80; VuTC01074), Photosystem II PsbX(P-II PsbX; VuTC14443), plasma membrane ATPase 4(VuTC29946), photosystem II stability/assembly factor HCF136 (VuTC01110) and 40S ribosomal protein S19-1(VuTC15418). The strong expression of HSC80 in floral shoot apices until six days post anthesis has been observed by Koning *et al*., (1992). Further, the expression of P-II PsbX(Shi et al., 1999) and plasma membrane ATPase 4(Zhang et al., 2017) in the flowers of *Spinacia oleracea* and *Arabidopsis thaliana* respectively has also been reported. Meurer *et al*., (1998) have demonstrated that the *Arabidopsis thaliana* plants with mutant HCF136 deficient in PSII activity were failed to produce flowers. Another transcript annotated as 40S ribosomal protein S19-1, was shown to be differentially expressed by Yan *et al*., (2014) while analyzing the differential expression in anther and stigma of *Eruca Sativa* during pre-bloom and after flowering stages. One transcript (VUTC21295) associated with both flowering and maturity, which codes for aldo-keto reductase has been discussed earlier regarding its association with the related traits. Its appearance within the top 10 expressed transcripts in all the three stages confirms its association. A transcript VuTC26129 annotated as WD repeat-containing protein 48(WDR48) and associated with maturity appeared within 10 expressed transcripts for all the three stages. A WDR48 analog in *Arabidopsis thaliana* has been demonstrated to interact and activate a deubiquitinase UBP3 (Baskerville et al., 2020). Additionally, plant deubiquitinases are reported to control flowering, embryogenesis, pollen and seed development (Schmitz et al., 2009; Doelling et al., 2001; Doelling et al., 2007). The transcript VuTC25355 associated with seed weight was annotated as 60S ribosomal protein L18a (RPL18a) and the role of RPL18a in embryo development has been established by Yan *et al,* (2016).

Apart from all the above discussed genes, few major genes like WRKY transcription factor 1(VuTC01132: seed weight), E3 ubiquitin-protein ligase RHG1A (VuTC02166: flowering), DEAD-box ATP-dependent RNA helicase 27(VuTC12393: seed weight), pentatricopeptide repeat-containing protein (VuTC09951: seed weight, VuTC31321: maturity) were found to contain trait associated markers. A *WRKY1* gene of *Solanum chacoense* has been reported be involved in the process of seed development having a specific role during the process of embryogenesis and was also found to be highly expressed in fertilized ovules at the late torpedo stage in wild potato (Lagacé and Matton, 2004). Shu and yang, (2017) have reported that the E3 ubiquitin-protein ligase RHG1A protein containing RING-H2_TTC3 type domain involved in the flowering time control and light response. Further, the role of deadbox RH27 in the process of seed development in *Arabidopsis thaliana* has been established by Hou *et al*. (2021) through a mutagenesis experiment. They reported that a recessive mutation in the gene produced shriveled or wrinkled seed.

The associative transcriptomics approach followed in the present investigation revealed a total of 127 markers on 87 transcripts associated with flowering, maturity and seed weight traits. Besides, 81 markers on 46 transcripts for flowering, 23 markers on 19 transcripts for maturity, 20 markers on 18 transcripts for seed weight, 2 markers on 2 transcripts for both seed weight and maturity, 1 marker on 1 transcript for both flowering and maturity were found associated. However, 1 transcript was found associated with both flowering and seed weight with different markers for both the traits. The functional annotation of transcripts revealed the corresponding gene description, domains, super families and GO terms of all the associated transcripts. Further, the role of the identified genes in influencing the corresponding traits has been corroborated with earlier findings in other crops. The association analysis involving SNPs obtained from transcriptome-based variant calling followed in this study is expected to provide deep insights into genetic mechanisms governing economically important production traits for various crops.

## Supplementary data

The following supplementary data are available at JXB online.

Table S1. The detailed available passport information on 100 considered accessions including date of collection, site of origin and cultivar name

Table S2. The number of bases and reads before and after filtration from the transcriptome data of 100 accessions of rice bean

Table S3. The chromosomal localization of markers with respect to the genome of the rice bean cultivar FF25

Table S4. The annotation of transcripts in terms of chromosomal localization and molecular function

Fig. S1. A hierarchical representation of GO terms for all the associated transcripts

## Author contributions

TKS: Data analysis and writing the original draft; SKV, NPS: Data analysis; Gayacharan, DCJ, SKV: Generation of phenotyping data; DPW, MS, RB, SKP, DCJ, GPS: writing, review and editing, AKS: conceptualization, writing, review and editing.

## Data availability

The data used in this study has been submitted to NCBI under the Bioproject accession PRJNA916051.

## References

Abraham G, Inouye M. 2014. Fast principal component analysis of large-scale genome-wide data. PLoS One 9(4), e93766. doi: doi.org/10.1371/journal.pone.0093766.

Andrews S. 2010. FastQC: A Quality Control Tool for High Throughput Sequence Data. http://www.bioinformatics.babraham.ac.uk/projects/fastqc/.

Baskerville A, Donahue J, Gillaspy G, et al. 2020. Identification of a WD-repeat protein that binds and activates the deubiquitinase UBP3 from *Arabidopsis thaliana*. BIOS 91(2), 90–99. https://doi.org/10.1893/BIOS-D-18-00029.

Bland JM, Altman DG.1995. Multiple significance tests: The Bonferroni method. BMJ 310(6973), 170. doi:10.1136/bmj.310.6973.170

Bolger AM, Lohse M, Usadel B. 2014.Trimmomatic: a flexible trimmer for Illumina sequence data. Bioinformatics 30(15), 2114–20. doi: 10.1093/bioinformatics/btu170.

Bowman AW, Azzalini A. 2021. R package ‘sm’: nonparametric smoothing methods (version 2.2-5.7). http://www.stats.gla.ac.uk/~adrian/sm.

Bradbury PJ, Zhang Z, Kroon DE, et al. 2007. TASSEL: Software for association mapping of complex traits in diverse samples. Bioinformatics 23, 2633–2635.

Cappetta E, Andolfo G, Guadagno A, et al. 2021. Tomato genomic prediction for good performance under high-temperature and identification of loci involved in thermotolerance response. Horticulture Research 8, 212. doi:10.1038/s41438-021-00647-3.

Chandel KP, Joshi BS, Arora RK, et al. 1978. Rice bean - a new pulse with high potential. Indian Farming 28, 19–22.

Cheng ZJ, Zhao XY, Shao XX, et al. 2014. Abscisic acid regulates early seed development in Arabidopsis by ABI5-mediated transcription of SHORT HYPOCOTYL UNDER BLUE1. Plant Cell 26(3), 1053–68. doi: 10.1105/tpc.113.121566.

Contreras S, Bennett MA, Metzger JD, et al. 2008. Maternal light environment during seed development affects lettuce seed weight, germinability, and storability. Hortscience 43, 845–852.

Dahiphale AV, Kumar S, Sharma N, et al. 2017. Rice bean-A Multipurpose, Underutilized, Potential Nutritive Fodder Legume - A Review. Journal of Pure and Applied Microbiology, 11 (1), 433–439. doi:10.22207/JPAM.11.1.57.

De Carvalho NM, Vieira RD.1996. Rice bean (*Vigna umbellata* (Thunb.) Ohwi et Ohasi). In: Nwokolo E, Smartt J, eds. Food and Feed from Legumes and Oilseeds. Boston: Springer, 222–228. doi:10.1007/978-1-4613-0433-3-25.

Dhillon PK, Tanwar B. 2018. Rice bean: A healthy and cost-effective alternative for crop and food diversity. Food Security. 10, 525–535. doi:10.1007/s12571-018-0803-6.

Dinesh A, Patil A, Zaidi PH, et al. 2016. Genetic diversity, linkage disequilibrium and population structure among CIMMYT maize inbred lines, selected for heat tolerance study. Maydica 61 (3), 1–7.

Doelling JH, Yan N, Kurepa J, et al. 2001. The ubiquitin-specific protease UBP14 is essential for early embryo development in *Arabidopsis thaliana*. The Plant Journal 27(5), 393–405.

Doelling JH, Phillips AR, Soyler-Ogretim G, et al. 2007. The ubiquitin-specific protease subfamily UBP3/ UBP4 is essential for pollen development and transmission in *Arabidopsis*. Plant Physiology 145, 801–813.

Dwivedi GK. 1996. Tolerance of some crops to soil acidity and response to liming. Journal of the Indian Society of Soil Science 44, 736–41.

Earl DA, vonHoldt BM. 2012. Structure Harvester: a website and program for visualizing STRUCTURE output and implementing the Evanno method. Conservation Genetics Resources 4 (2), 359–36. doi: 10.1007/s12686-011-9548-7.

Finkelstein RR, Gampala SS, Rock CD. 2002. Abscisic acid signaling in seeds and seedlings. Plant Cell 14(Suppl), S15–45. doi: 10.1105/tpc.010441.

Gautam R, Kumar N, Yadavendra JP, et al. 2007. Food security through rice bean research in India and Nepal (FOSRIN). Report 1. Distribution of rice bean in India and Nepal. Local Initiatives for Biodiversity, Research and Development, Pokhara, Nepal and CAZS Natural Resources, College of Natural Sciences, Bangor University, Wales, UK.

González AM, Yuste-Lisbona FJ, Saburido S, et al. 2016. Major contribution of flowering time and vegetative growth to plant production in common bean as deduced from a comparative genetic mapping. Frontiers in Plant Science 7,1940. doi: 10.3389/fpls.2016.01940.

Gu W, Zhu J, Wallace DH, et al. 1998. Analysis of genes controlling photoperiod sensitivity in common bean using DNA markers. Euphytica 102, 125–132. doi:10.1023/A:1018340514388.

Guan J, Zhang J, Gong D. et al. 2022. Genomic analyses of rice bean landraces reveal adaptation and yield related loci to accelerate breeding. Nature Communications 13, 5707. doi: 10.1038/s41467-022-33515-2.

Gupta PK, Kulwalb PL, Jaiswal V. 2019. Association mapping in plants in the post-GWAS genomics era. Advances in Genetics doi:10.1016/bs.adgen.2018.12.00.

Hao Y, Zong X, Ren P, et al. 2021. Basic Helix-Loop-Helix (bHLH) Transcription Factors Regulate a Wide Range of Functions in Arabidopsis. International Journal of Molecular Sciences 22(13), 7152. doi: 10.3390/ijms22137152.

Hong SE, Kneissl J, Cho A, et al. 2022. Transcriptome-based variant calling and aberrant mRNA discovery enhance diagnostic efficiency for neuromuscular diseases. Journal of Medical Genetics 59(11), 1075–1081. doi: 10.1136/jmedgenet-2021-108307.

Hou XL, Chen WQ, Hou Y, et al. 2021.DEAD-BOX RNA HELICASE 27 regulates microRNA biogenesis, zygote division, and stem cell homeostasis. Plant Cell m33(1), 66–84. doi: 10.1093/plcell/koaa001.

Huang M, Liu X, Zhou Y, et al. 2019.BLINK: a package for the next level of genome-wide association studies with both individuals and markers in the millions, GigaScience 8(2), giy154. doi:10.1093/gigascience/giy154.

Isemura T, Kaga A, Tomooka N, et al. 2010. The genetics of domestication of rice bean, *Vigna umbellata*. Annals of botany 106(6), 927–944.

Ishikawa T, Maruta T, Yoshimura K, et al. 2018. Biosynthesis and Regulation of Ascorbic Acid in Plants. In: Gupta D, Palma J, Corpas F, eds. Antioxidants and Antioxidant Enzymes in Higher Plants. Cham: Springer, doi:10.1007/978-3-319-75088-0_8.

Jehl F, Degalez F, Bernard M, et al. 2021. RNA-Seq Data for Reliable SNP Detection and Genotype Calling: Interest for Coding Variant Characterization and Cis-Regulation Analysis by Allele-Specific Expression in Livestock Species. Frontiers in Genetics 12, 655707. doi: 10.3389/fgene.2021.655707.

Kanno Y, Jikumaru Y, Hanada A, et al. 2010. Comprehensive hormone profiling in developing Arabidopsis seeds: Examination of the site of ABA biosynthesis, ABA transport and hormone interactions. Plant Cell Physiology 51, 1988–2001.

Kaul T, Easwaran M, Thangaraj A, et al. 2022. De novo genome assembly of rice bean (Vigna umbellata) – A nominated nutritionally rich future crop reveals novel insights into flowering potential, habit, and palatability centric – traits for efficient domestication. Frontiers in Plant Science 13, 739654. doi: 10.3389/fpls.2022.739654.

Koning AJ, Rose R, Comai L. 1992. Developmental expression of tomato heat-shock cognate protein 80. Plant Physiology 100(2), 801–811, doi: 10.1104/pp.100.2.801.

Kotchoni SO, Larrimore KE, Mukherjee M, et al. 2009. Alterations in the endogenous ascorbic acid content affect flowering time in Arabidopsis. Plant Physiology 149(2), 803–15. doi: 10.1104/pp.108.132324.

Kwak M, Velasco D, Gepts P. 2008. Mapping Homologous Sequences for Determinacy and Photoperiod Sensitivity in Common Bean (*Phaseolus vulgaris*), Journal of Heredity 99(3), 283–291. https://doi.org/10.1093/jhered/esn005.

Lagacé M, Matton DP. 2004. Characterization of a WRKY transcription factor expressed in late torpedo-stage embryos of Solanum chacoense. Planta 219, 185–189. doi:10.1007/s00425-004-1253-2.

Li H, Durbin R. 2009. Fast and accurate short read alignment with Burrows–Wheeler transform. Bioinformatics 25(14), 1754–1760.

Li H, Handsaker B, Wysoker A, et al. 2009. The Sequence Alignment/Map format and SAMtools. Bioinformatics 25(16), 2078–9. doi: 10.1093/bioinformatics/btp352.

Lipka AE, Tian F, Wang Q, et al. 2012. GAPIT: genome association and prediction integrated tool. Bioinformatics 28(18), 2397–2399, doi:10.1093/bioinformatics/bts444.

Liu X, Huang M, Fan B, et al. 2016. Iterative Usage of Fixed and Random Effect Models for Powerful and Efficient Genome-Wide Association Studies. PLoS Genetics 12(2), e1005767. doi:10.1371/journal.pgen.1005767.

Lonardi S, Muñoz-Amatriaín M, Liang Q, et al. 2019. The genome of cowpea (Vigna unguiculata [L.] Walp.). Plant Journal 98(5),767–782. doi: 10.1111/tpj.14349.

Marciniak K, Przedniczek K. 2021. Anther dehiscence is regulated by gibberellic acid in yellow lupine (*Lupinus luteus* L.). BMC Plant Biology 21(1), 314. doi: 10.1186/s12870-021-03085-4.

McCarty DR.1995.Genetic control and integration of maturation and germination pathways in seed development, Annual Review of Plant Physiology and Plant Molecular Biology 46, 71–93.

Meurer J, Plücken H, Kowallik KV, et al. 1998. A nuclear-encoded protein of prokaryotic origin is essential for the stability of photosystem II in *Arabidopsis thaliana*. The EMBO Journal 17(18), 5286–5297. doi: 10.1093/emboj/17.18.5286.

Mohan VR, Janardhanan K. 1994. Chemical and nutritional evaluation of two germplasms of the tribal pulse, Bauhinia racemosa Lamk. Plant Foods for Human Nutrition 46(4), 367–74. doi: 10.1007/BF01088438.

NAS. 1979. Tropical Legumes: Resources for the Future. National Academy of Science, Report of an Ad Hoc Panel of the advisory committee on technology innovation. https://nap.nationalacademies.org/read/19836/chapter/1#iii.

Ni M, Tepperman JM, Quail PH. 1998. PIF3, a phytochrome-interacting factor necessary for normal photoinduced signal transduction, is a novel basic helix-loop-helix protein. Cell 95(5), 657–67. doi: 10.1016/s0092-8674(00)81636-0.

Pattanayak A, Roy S, Sood S, et al. 2019. Rice bean: a lesser known pulse with well-recognized potential. Planta 250(3), 873–890.

Penning TM. 2015. The aldo-keto reductases (AKRs): Overview. Chemico-Biological Interactions 234, 236–46. doi: 10.1016/j.cbi.2014.09.024.

Philippe F, Pelloux J, Rayon C. 2017. Plant pectin acetylesterase structure and function: new insights from bioinformatic analysis. BMC Genomics 18, 456. doi:10.1186/s12864-017-3833-0

Price A, Patterson N, Plenge R, et al. 2006. Principal components analysis corrects for stratification in genome-wide association studies. Nature Genetics 38, 904–909.

Pritchard JK, Stephens M, Donnelly P.2000. Inference of population structure using multilocus genotype data. Genetics 155, 945–959.

Schmid M, Davison T, Henz S, et al. 2005. A gene expression map of *Arabidopsis thaliana* development. Nature Genetics 37, 501–506. doi:10.1038/ng1543.

Schmitz RJ, Tamada Y, Doyle MR, et al. 2009. Histone H2B deubiquitination is required for transcriptional activation of FLOWERING LOCUS C and for proper control of flowering in Arabidopsis. Plant Physiology 149, 1196–1204.

Shi LX, Kim SJ, Marchant A, et al. 1999.Characterisation of the PsbX protein from Photosystem II and light regulation of its gene expression in higher plants. Plant Molecular Biology 40(4),737–744. doi: 10.1023/a:1006286706708.

Shu K, Yang W. 2017. E3 Ubiquitin Ligases: Ubiquitous Actors in Plant Development and Abiotic Stress Responses. Plant Cell Physiology 58(9), 1461–1476. doi: 10.1093/pcp/pcx071.

Singh D, Singh CK, Taunk J, et al. 2019. Genome wide transcriptome analysis reveals vital role of heat responsive genes in regulatory mechanisms of lentil (Lens culinaris Medikus). Scientific Reports 9(1), 12976. doi: 10.1038/s41598-019-49496-0.

Smil V. 1997. Some unorthodox perspectives on agricultural biodiversity. The case of legume cultivation. Agriculture, Ecosystems & Environment 62, 135–144.

Somta P, Kaga A, Tomooka N, et al. 2006. Development of an interspecific Vigna linkage map between Vigna umbellata (Thunb.) Ohwi & Ohashi and V. nakashimae (Ohwi) Ohwi & Ohashi and its use in analysis of bruchid resistance and comparative genomics. Plant Breeding 125, 77–84. doi:10.1111/j.1439-0523.2006.01123.x.

Tamba CL, Ni YL, Zhang YM. 2017. Iterative sure independence screening EM-Bayesian LASSO algorithm for multi-locus genome-wide association studies. PLoS Computational Biology 13(1), e1005357. doi: 10.1371/journal.pcbi.1005357.

Tamba CL, Zhang YM. 2018. A fast mrMLM algorithm for multi-locus genome-wide association studies, bioRxiv doi:10.1101/341784.

Tian J, Isemura T, Kaga A, et al. 2013. Genetic Diversity of the rice bean (*Vigna Umbellata*) Genepool as Assessed by SSR Markers. Genome 56, 717–727. doi:10.1139/gen-2013-0118.

Tripathi A, Iswarya V, Singh N et al. 2021. Chapter 4 - Chemistry of pulses—micronutrients, In: Tiwari BK, Gowen A, McKenna B, eds. Pulse Foods (Second Edition). Cambridge: Academic Press, 61–86. doi:10.1016/B978-0-12-818184-3.00004-0.

Van Eldik GJ, Wingens M, Ruiter RK et al. 1996. Molecular analysis of a pistil-specific gene expressed in the stigma and stylar cortex of *Solanum tuberosum*. Plant Molecular Biology 30, 171–176. doi: 10.1007/BF00017811.

Voorrips RE. 2002. MapChart: Software for the graphical presentation of linkage maps and QTLs. The Journal of Heredity 93 (1),77–78.

Wang SB, Feng JY, Ren WL, et al. 2016. Improving power and accuracy of genome-wide association studies via a multi-locus mixed linear model methodology. Scientific Reports 6, 19444. doi:10.1038/srep19444.

Wen YJ, Zhang H, Ni YL, et al. 2018. Methodological implementation of mixed linear models in multi-locus genome-wide association studies. Briefings in Bioinformatics 19(4), doi: 10.1093/bib/bbw145.

Xiao Y, Liu H, Wu L, et al. 2017. Genome-wide association studies in Maize: praise and stargaze. Molecular Plant 10, 359–374.

Yamamoto Y, Aminaka R, Yoshioka M, et al. 2008. Quality control of photosystem II: impact of light and heat stresses. Photosynthesis Research 98, 589–608. doi.org/10.1007/s11120-008-9372-4.

Yan F, Wan-cang S, Jun-yan W, et al. 2014. Differential display and expression analysis of self-compatibility associated gene in *Eruca sativa*. Chinese Journal of Oil Crop Sciences, 36(5), 580–585.

Yan H, Chen D, Wang Y, et al. 2016. Ribosomal protein L18aB is required for both male gametophyte function and embryo development in *Arabidopsis*. Scientific Reports 6, 31195. https://doi.org/10.1038/srep31195.

Yu H, Deng Z, Xiang C, et al. 2014. Analysis of diversity and linkage disequilibrium mapping of agronomic traits on B-genome of wheat. Journal of Genomics 2, 20–30. doi: 10.7150/jgen.4089.

Yu J, Pressoir G, Briggs WH, et al. 2006. A unified mixed-model method for association mapping that accounts for multiple levels of relatedness. Nature Genetics 38, 203–208. doi: 10.1038/ng1702.

Yu Y, Ouyang Y, Yao W. 2018. shinyCircos: an R/Shiny application for interactive creation of Circos plot. Bioinformatics 34(7), 1229–1231. doi: 10.1093/bioinformatics/btx763.

Zhang J, Wei J, Li D, et al. 2017. The Role of the Plasma Membrane H+-ATPase in Plant Responses to Aluminum Toxicity. Frontiers in Plant Science 8, 1757. doi: 10.3389/fpls.2017.01757.

Zhang YM, Jia Z, Dunwell JM. 2019. Editorial: The Applications of New Multi-Locus GWAS Methodologies in the Genetic Dissection of Complex Traits. Frontiers in Plant Science 10, 100. doi: 10.3389/fpls.2019.00100.

Zhao K, Tung CW, Eizenga G, et al. 2011. Genome-wide association mapping reveals a rich genetic architecture of complex traits in Oryza sativa. Nature Communications 2, 467. doi:10.1038/ncomms1467.

Zhou H, Yin H, Chen J, et al. 2016. Gene-expression profile of developing pollen tube of Pyrus bretschneideri. Gene Expression Patterns 20(1), 11–21. doi: 10.1016/j.gep.2015.10.004.

